# A DNA-based T cell receptor reveals a role for receptor clustering in ligand discrimination

**DOI:** 10.1101/062877

**Authors:** Marcus J. Taylor, Kabir Husain, Zev J. Gartner, Satyajit Mayor, Ronald D. Vale

**Affiliations:** Dept. of Cellular and Molecular Pharmacology, University of California San Francisco, USA; Howard Hughes Medical Institute, University of California San Francisco, USA; National Centre for Biological Sciences, Bangalore, India; The Simons Centre for the Study of Living Machines, Bangalore, India; HHMI Summer Institute, Woods Hole MA; Dept. of Pharmaceutical Chemistry, University of California San Francisco, USA

## Abstract

T cells mount an immune response by measuring the binding strength of its T cell receptor (TCR) for peptide-loaded MHCs (pMHC) on an antigen-presenting cell. How T cells convert the lifetime of the extracellular TCR-pMHC interaction into an intracellular signal remains unknown. Here, we developed a synthetic signaling system in which the extracellular domains of the TCR and pMHC were replaced with short hybridizing strands of DNA. Remarkably, T cells can discriminate between DNA ligands differing by a single base pair. Single molecule imaging reveals that signaling is initiated when single ligand-bound receptors are converted into clusters, a time-dependent process requiring ligands with longer bound times. A computation model reveals that receptor clustering serves a kinetic proofreading function, enabling ligands with longer bound times to have disproportionally greater signaling outputs. These results suggest that spatial reorganization of receptors plays an important role in ligand discrimination in T cell signaling.

## Introduction

The recognition of foreign antigens by T cells begins with a binding interaction between cell surface peptide-loaded MHC (pMHC) and the T cell receptor (TCR) expressed on the surface of T cells. A pMHC-TCR interaction of sufficient strength triggers the phosphorylation of immunoreceptor tyrosine activation motifs in the TCRζ and associated CD3 chains by the Src family kinase Lck^1^. The mechanism by which pMHC engagement leads to TCR phosphorylation remains controversial; current models include receptor conformational changes^2^ and exclusion of the inhibitory transmembrane phosphatase CD45 from zones of pMHC-TCR engagement^3^. The phosphorylated ITAM domains then recruit the kinase ZAP70 ^4^, which in turn phosphorylates the adapter protein LAT (Linker for Activation of T cells)^5^. Downstream of LAT, numerous signaling pathways become activated, including the MAP kinase pathway, actin polymerization^6^, elevation of intracellular calcium, and large scale changes in transcription^7-9^.

A remarkable feature of the T cell signaling is its ability to respond to and clear the body of viral and microbial infection but not mount a destructive response to the body’s own cells. Through genetic recombination, each T cell expresses a unique TCR with its own binding specificity^10^. Unlike many cell surface receptors that interact with a single or limited number of ligands, the TCR is presented with an immense number of different peptides loaded onto MHC molecules. The vast majority of these peptides are low affinity antigens derived from the body’s own cells. In order not to generate a harmful auto-immune response, mature T cells must ignore the majority of these low affinity interactions and selectively activate in response to pMHC loaded with higher affinity foreign peptides.

Previous studies have implicated the lifetime of the TCR-pMHC interaction as a key determinant between activating and non-activating pMHC molecules ^11-13^. Remarkably, even a few-fold variation in the off-rates of different peptide-bound MHCs for a given TCR can result in an all-or-none differences in downstream signaling outputs at physiological ligand densities ^14^. However, a mechanistic explanation of how lifetime of an extracellular interaction is ‘read out’ and then converted to an intracellular signal is not well understood. A theory of “kinetic proofreading” was developed to explain how relatively small differences in receptor bound time might be discriminated and lead to more binary downstream outputs ^15^. In the general formulation of kinetic proofreading, signaling is triggered by a linked set of reactions that require the continuous occupancy of the ligand-receptor complex; if the ligand dissociates, then these reactions are rapidly reversed and the receptor is reset to a inactive state. Many different molecular mechanisms have been put forth for the kinetic proofreading steps, including enzymatic reactions (e.g. phosphorylation)^15-17^, receptor conformational changes ^18,19^ and receptor dimerization ^20^. However, compelling evidence for kinetic proofreading is lacking, and it also remains controversial whether kinetic proofreading begins at the level of the receptor ^14^ or farther downstream ^21^.

Taking a reductionist approach to understand T cell signaling and ligand discrimination, we sought to engineer a T cell signaling system in which receptor-ligand affinity can be precisely tuned over a wide dynamic range and the receptor behavior and signaling output measured at the single molecule level and without influence from other co-receptors ^22^. Previous work has shown that the extracellular ligand binding regions of the TCR could be replaced with a single chain antibody, which upon binding to its antigen on another cell membrane, will initiate T cell signaling and activation ^23,24^. Currently, T cells expressing such chimeric antigen receptors (CARs) are being tested for their ability to eliminate cancer cells ^25^. Based upon the work of CARs, we reasoned that the extracellular domains of the TCR and pMHC could be replaced by complementary strands of DNA and that DNA hybridization might act as receptor-ligand interaction. The advantage of using DNA is that its nucleotide composition can be varied to provide exquisite and predictable control of the strength of the ligand-receptor interaction. Using this system and single molecule live cell imaging, we have found a time-dependent conversation of single receptor-ligands into receptor clusters is required for efficient phosphorylation of TCR phosphorylation and the recruitment of ZAP70. The formation of receptor clusters arises from a dramatic increase in the ligand binding on-rate adjacent to pre-existing ligated receptors. These results, in combination with mathematical modeling, reveal that the spatial organization of receptor-ligand complexes provides an early proofreading step in T cell signaling.

## Results

### Development and characterization of a DNA – Chimeric Antigen Receptor

We created a nucleic acid-based synthetic DNA-CARζ that consists of a ssDNA covalently reacted to an extracellular SNAP tag protein ^14,26^ that was fused to a transmembrane domain and intracellular CD3ζ chain (Fig. 1A). To avoid any potential signaling cross-talk with the native receptor, we expressed this DNA-CARζ in a TCR negative Jurkat cell line (JRT3)^27^. To stimulate the DNA-CARζ, we replaced the antigen-presenting cell with a planar supported lipid bilayer (SLB) ^28-30^ functionalized with a freely diffusing CLIP protein covalently bound with a complimentary strand of ssDNA (Methods). A single fluorescent dye also could be incorporated into the ligand DNA-CLIP complex in a non-perturbing and stoichiometric manner, allowing single molecule observations. T cells and APCs initially interact through adhesion molecules (e.g. ICAM-LFA1) or other co-receptors, which also have signaling functions ^31^. To enable our DNA-CARζ T cells to adhere to the SLB without any co-stimulus, we made use of a synthetic DNA ‘adhesion system’ that de-couples adhesion from cell signaling^32^ (Fig. 1A).

**Fig. 1.**
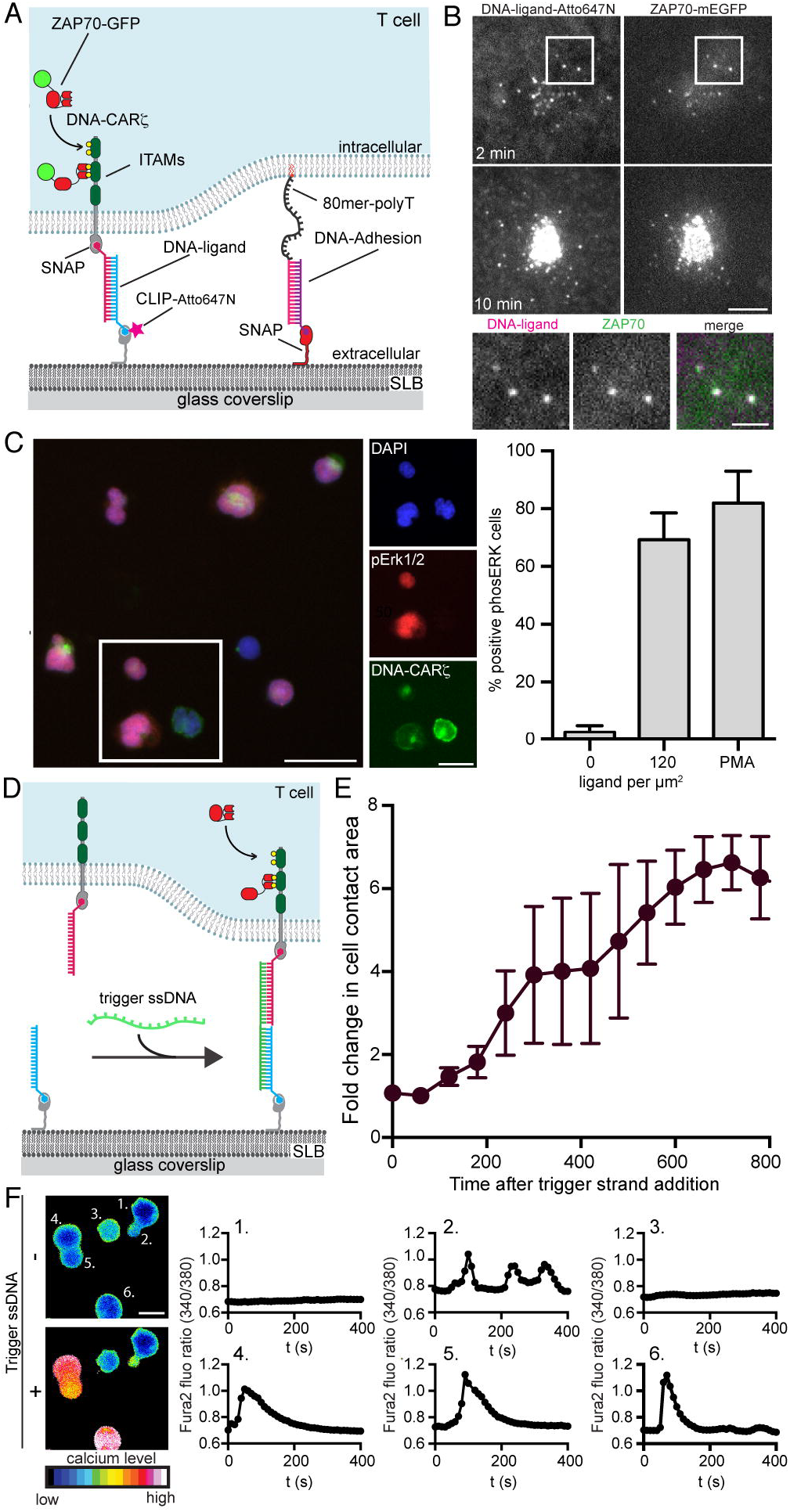
A DNA-CARζ capable of triggering T cell signaling. **A.** Schematic of the DNA-based chimeric antigen receptor system (DNA-CARζ). The SNAPf tag and His10-CLIPf were covalently labeled with complementary strands of benzyl-guanine-DNA and benzyl-cytosine DNA respectively. Longer orthogonal DNA strands without transmembrane coupling to an intracellular signaling domain were used to adhere cells to the bilayer without inducing signaling. **B.** TIRF microscopy images of a JRT3 Jurkat cell expressing ZAP70-GFP and DNA-CARζ labeled with 16mer ssDNA after landing on a SLB with a complementary 16mer strand (120 molecules per µm^2^). Microclusters of ligand-receptor complexes formed within ~2 min and recruited ZAP70-GFP (inset) and then moved centripetally and coalesced near the cell center at 10 min. Scale bar, 5 µm; inset scale bar, 2 µm. **C.** To measure activation of the MAP kinase pathway, cells (15 min after SLB contact) were stained for phosphoERK (red); DAPI staining of nuclei (blue) and the DNA-CARζ (green). Bar, 50 µm. Insert shows higher magnification of three cells. Inset bar, 20 µm. Quantification (see Fig. S2, Methods) of the MAP kinase pathway activation by the 16mer DNA ligand compared to PMA (10 ng/ml). Mean ± s.d. of 6 experiments (>2500 cells scored per experiment). **D.** A schematic of a triggerable DNA-CARζ designed to examine temporal responses of T cell signaling. Although included in experiments, the adhesion strand shown in panel **a** is omitted for clarity. **E.** Cell spreading on the SLB as a function of time after adding the DNA trigger strand. Plotted is average fold change in cell area as a function of time after addition of trigger strand. Mean ± s.e.m. from 3 separate experiments (3-7 cells analyzed per experiment). **F.** Pseudo colour image of calcium level using the ratiometric Fura-2 calcium dye. Plots of the 340 nm/380 nm Fura-2 emission ratio showing the change in intracellular calcium levels from the six cells after adding the DNA trigger strand. Bar, 10 µm.

A high affinity pMHC-TCR leads to increased intracellular calcium, MAP kinase activation, re-organization of the actin cytoskeleton and the re-localization of transcription factors to the nucleus ^7,33,34^. To assess whether the DNA-based CAR is capable of transmitting similar intracellular signals upon ligand binding, we first tested a high affinity 16mer DNA base pair interaction (predicted off-rate of >7 hr, as estimated from computational analysis ^35^). When these cells were plated on SLBs with a high ligand density of ~120 molecules per µm^2^, we observed the rapid reorganization of ligand-bound receptors into submicron clusters that recruited the tyrosine kinase ZAP70-GFP(Fig. 1B; Movie S1). These clusters were dynamic and translocated centripetally from the periphery to the cell center (Movie S1). This receptor behavior is similar to that reported for antibody or pMHC activated TCR ^28,29,36^. The majority (~65%) of the DNA-CARζ T cells also signaled through the MAP kinase cascade, as indicated by strong phospho-ERK (pERK) staining of the nucleus (Fig. 1C); this response was comparable to that produced by the strong stimulus of phorbol myristate acetate (PMA) (Fig. 1C) and to that reported for TCR-pMHC in native T cells^37^.

To examine the kinetics of signaling, we designed a system in which cells could be triggered to signal in a synchronous manner after they were adhered to the SLB through the inert DNA-adhesion system. To achieve such temporal control, we designed non-complementary ssDNA for the ligand and receptor and then introduced an oligonucleotide that could hybridize to both receptor and ligand DNA and thus bridge the two together (Fig. 1D). Following the addition of this “trigger strand”, the adhered cells spread rapidly (2-5 min), a result of the activation of actin polymerization (Fig. 1E and Fig. S1B). Intracellular calcium also rose after ~1 min (Fig. 1F; Movie S2), and CD69, a TCR activation marker, was expressed on the cell surface at 4 hr (Fig. S1D). The timing of these responses are similar to those reported previously ^23^. Thus, the DNA-based CARζ induces similar intracellular signaling responses to those described for T cells triggered through TCR and pMHC.

### Ligand discrimination by the DNA – CARζ

Next, we tested how signaling is influenced by the length and sequence of the DNA using automated microscopy and image analysis of pERK staining as a readout (Fig. S2A,B)^38^. For these experiments, we used the direct hybridizing ligand-receptor pair (Fig. 1A), since the ligand concentrations on the SLB can be carefully controlled and varied (Methods). Compared to the 16mer oligo, ligand dose response curves of the 13mer and 12mer oligo ligands produced progressively weaker MAP kinase signaling (Fig. 2A, B). The 11mer did not elicit a measurable pERK response above background, even at the highest ligand density. We then attempted to restore signaling to the 11mer by mutating adenine/thymine (A/T) base pairs to guanine/cytosine (G/C) base pairs, which increases the binding free energy of hybridization (Table S1). Remarkably, a single A/T to G/C mutation converted the initial non-signaling 11mer receptor to one that could elicit a pERK response at high ligand densities, and each additional G/C mutation increased the potency of the DNA receptor (Fig. 2C).

**Fig. 2.**
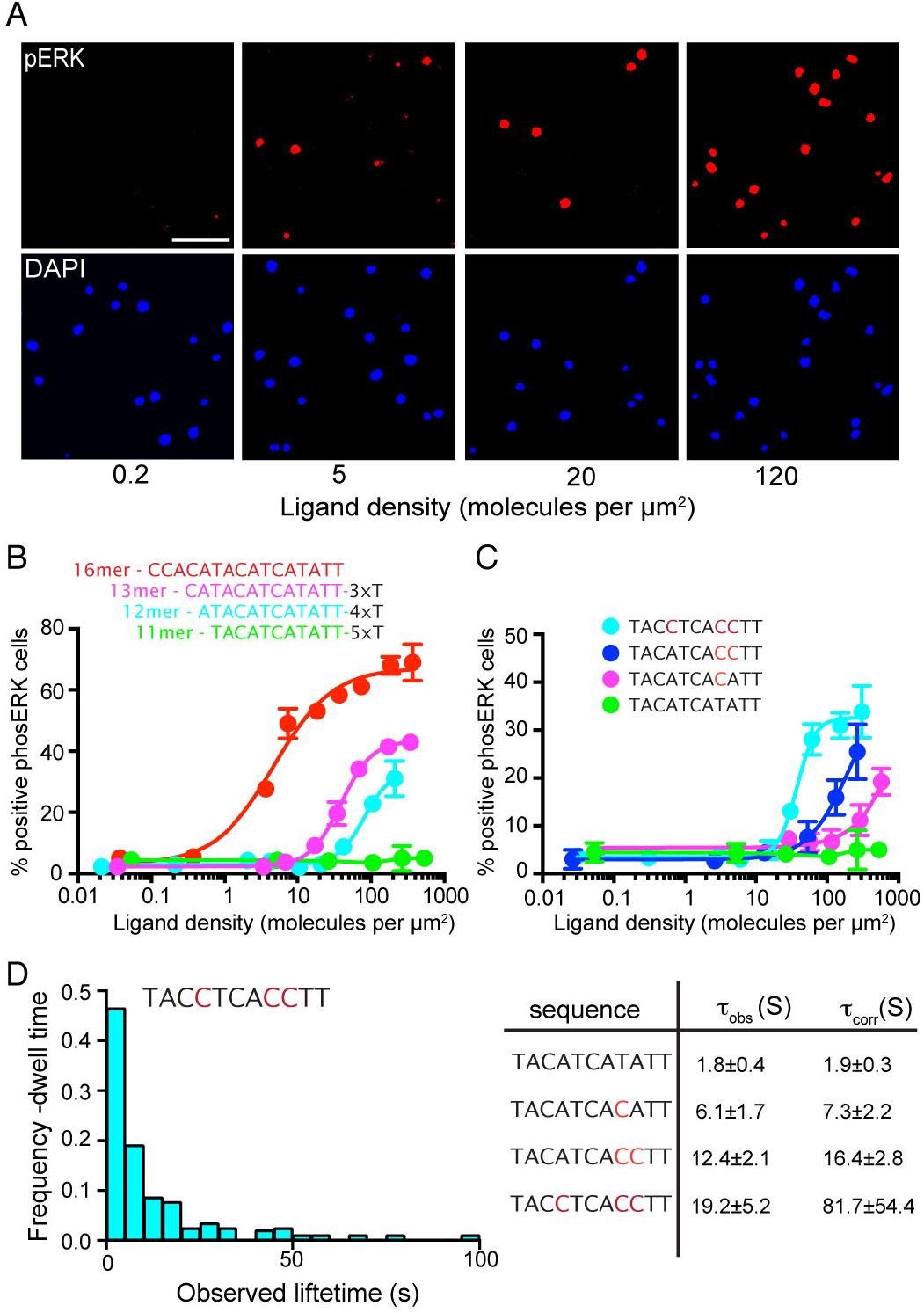
Modulation of T cell activation by DNA-CARζ length and sequence. **A.** PhophoERK staining of JRT3 cells responding to an increasing density of 16mer DNA ligand on SLBs (15 min). Threshold images are shown corresponding to the phosphERK staining and DAPI channel. Scale bar, 100 µm. **B.** Dose response curves for DNA ligands of varying hybridization length. The total ligand DNA length remained constant by adding nonhybridizing Ts. Mean ± s.d. (n = 3). **C.** Stepwise conversion of A/T to G/C base pairs increases the potency of the 11mer DNA ligand for inducing phosphoERK signaling. Scale bar, 100 µm. Each data point represents the mean from one experiment, where each ligand density was measured in triplicate. Mean ± s.d. (n = 3). **D.** Measurement of the lifetime of single ligand-receptor interactions by TIRF microscopy. An example distribution of lifetimes from one DNA ligand. The observed (τ_obs_) and photobleach-corrected (τ_corr_) lifetimes are shown for the 11mers (mean ± s.e.m. from 3 separate experiments reported; for data of 13mer and 16mer and all histograms see Fig. S2).

Since theoretical models ^15,39,40^ and biochemical data ^12,13^ have suggested that the bound time of the TCR and pMHC plays a critical role in signaling, we directly measured the lifetime of individual DNA receptor-ligand interactions at the cell-SLB interface using single molecule total internal reflection fluorescence microscopy (TIRFM)^21^ (Fig. 2D, and Fig. S2). The bound time of the ligand for the receptor displayed a roughly exponential distribution (Fig. 2D and Fig. S2E, F), with an observed half-life of ~2 s for the non-activating 11mer and a half-life of ~19 s for the 11mer with three additional G/C bases (Fig. 2D). The much slower off-rate of the 16mer DNA ligand (predicted to be hours) could not be determined accurately, since these measurements were limited by the rate of photobleaching (Fig. S2E). Overall, the ligand bound times and the ligand densities on the SLB required for half-maximal responses are similar to those reported for TCR-pMHC in comparable bilayer activation experiments ^21^. Collectively, these results clearly show that increasing the GC base pairing of the receptor results in a longer ligand-receptor interaction and that the T cell signaling response can distinguish between DNA receptor-ligand interactions with small difference in binding free energy (~1 kcal/mol, Table S1).

### Single molecule imaging of DNA-CARζ phosphorylation

We next wanted to examine how the extracellular receptor-ligand bound time is translated into intracellular biochemistry, the first step being the phosphorylation of the ITAM domains of the DNA-CARζ receptor. Phosphorylation of the ITAMs leads to the recruitment of the kinase ZAP70 ^41^ which phosphorylates downstream targets. To measure phosphorylation of DNA-CARζ in live cells in real time, we examined the recruitment of ZAP70-GFP from the cytosol to receptor-ligand complexes by TIRFM (Fig. 3; Fig. S3E, F and Fig. S4). We first used the 16mer at a ligand density of 0.1 molecules per µm^2^, a density below the threshold required to elicit a pERK response. Under these conditions, single receptor-ligand binding interactions can be observed clearly (Fig. 3A; Fig. S4A; Movie S3). However, surprisingly only ~6% of the ligand binding events (trackable for 30 s or more) led to detectable ZAP70-GFP recruitment, despite the long bound time of the ligand-receptor interaction (Fig. 3B). In these rare cases, ZAP70-GFP recruitment was transient and lasted less than ~20 sec (Fig. 3C); this single molecule measurement reflects the off-rate of ZAP70 since the 16mer has an off-rate on the hour time scale. In this experiment, nonfluorescent endogenous ZAP70 could potentially outcompete ZAP70-GFP for binding to the ligated receptor, thus affecting measurements and conclusions. Thus, we repeated these experiments in the P116 ZAP70 null Jurkat cell line ^42^; very similar results were observed (Fig. S4B-D), confirming that ZAP70 is not efficiently recruited to single receptor-ligand complexes at low ligand densities. These results indicate that a long-lived ligand binding interaction *per se* is not sufficient to trigger receptor phosphorylation.

**Fig. 3.**
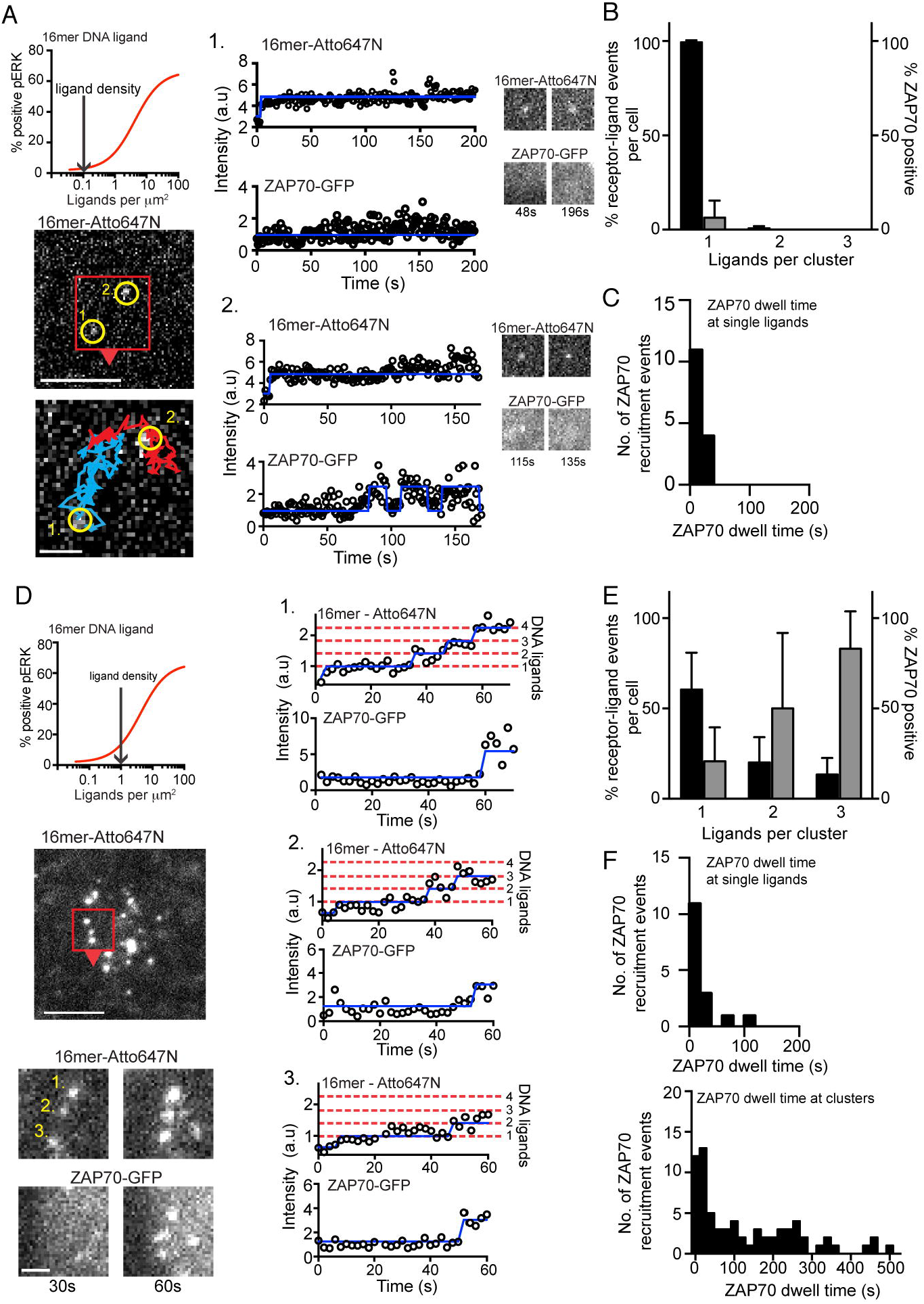
Receptor clustering increases the probability of ZAP70 recruitment. Measuring ligand and ZAP70-GFP binding to DNA-CARζ by TIRF imaging with a 16mer DNA ligand at 0.1 (a-c) and 1 ligand per µm^2^ (d-e). Dose response curves shown in panel a and d are based on data shown in Fig. 2b. The blue line overlaid the Atto647N and ZAP70-GFP fluorescent intensity represents detected step changes in fluorescence intensity marking new ligand binding events or the recruitment of ZAP70. The dashed red lines mark the quantal ligand fluorescent intensities for a given number of DNA ligands, (determined using a hidden Markov model analysis; see Fig. S3 and Supplemental Information). Single or multiple ligand binding events that persisted and could be followed for >30 s were scored for whether they recruited ZAP70-GFP. The initial point of ZAP70-GFP recruitment was referenced to the number of ligands within the receptor-ligand spot as in panel a and d. **A.** TIRF images of 16mer DNA ligands labeled with Atto647N. Single bound ligands are marked by yellow circles in region of interest (red box and arrowhead). Bar, 5 µm. Right panel, region of interest overlaid with the tracked single molecule trajectories. ROI bar, 1 µm. Right panels, the fluorescent intensity time series for the 16mer ligand and the corresponding ZAP70-GFP fluorescent intensity. Ligand 1 in (a) shows a typical example of a bound ligand-receptor pair that does not recruit a ZAP70-GFP. Ligand 2 in (**a**) shows a less common example of ZAP70-GFP recruitment (often transient as shown here) to a single bound 16mer ligand. **B.** Quantification of ZAP70-GFP recruitment at 0.1 ligand/µm^2^ of 16mer DNA ligand. Bar plot shows the number single bound ligand-receptors and clusters (percent of total, black bars) and ZAP70 recruitment to ligand-receptors (percent of total that recruit ZAP70, grey bars). 6.3 ± 9% of single bound ligands that persist greater than 30 s recruit ZAP70. Results are mean ± s.d. from n = 9 cells. **C.** Quantification of ZAP70 dwell time at single bound ligands of 16mer DNA ligand (n=15). **D.** TIRF images of 16mer DNA ligands at 1 ligand/µm^2^ (which elicits pERK signaling in ~20% of cells). Bar, 5 µm. Region of interest (red box and arrowhead) shows three ligand-receptor clusters (labeled 1-3). ROI bar, 1 µm. Fluorescent intensity time series from the DNA and ZAP70-GFP channels of the same 3 microclusters is shown on the right. For additional traces at various ligand densities, see Fig. S4 and Movies S3-6. **E.** Quantification of ZAP70 recruitment at 1 ligand/µm^2^ of 16mer DNA ligand. Organization of bar plot same as shown (b). 83 ± 20% or receptor-ligand clusters composed of 3 or more ligands recruit ZAP70 versus 20 ± 18% of single bound receptors. Results are mean ± s.d. from n = 6 cells. **F.** Distribution of ZAP70 dwell times at single bound receptors (n=16) and receptor-ligand clusters (n=70). For 32 of the clusters with ZAP70 dwells time of >100s, the measurement was truncated by the end of image acquisition.

Since binding of the high-affinity 16mer ligand to a single receptor did not lead to stable receptor phosphorylation, we next wished to understand what additional events might be needed to initiate this first step in signaling. To answer this question, we performed single molecules studies at a 10-fold higher 16mer ligand density (16mer ligand density of 1 molecule per µm^2^), a regime in which 20% of the cells generate a pERK response (Fig. 3D) and the ligand density was still low enough to enable clear single molecule imaging. At 1 molecule per µm^2^, single 16mer receptor-ligand pairs formed initially and then grew into small clusters on the membrane within a few minutes of cell contact with the SLB (Fig. 3D and Fig. S4E; Movie S4). By quantitating the fluorescence increase of these small receptor ligand clusters, we could estimate how many bound ligands were present in a cluster at the initial moment of ZAP70-GFP recruitment (Fig. 3D, Fig. S3). This analysis revealed that ZAP70-GFP is recruited more efficiently to clusters containing three or more ligated receptors (~80%) compared with single ligated receptors (~20%) (Fig. 3E). Furthermore, the bound time of ZAP70-GFP on a single ligated receptor was short (half-life of ~10 s, Fig. 3F and example 4 in Fig. S4E), as reported for at the lower ligand density on the bilayer (Fig. 3C). In striking contrast, nearly half of ZAP70-GFP molecules (48%) remained associated for 100 s or more with small receptor clusters (Fig. 3F). Thus, clusters composed of a just few bound receptors become phosphorylated and stably recruit ZAP70-GFP, whereas single ligated receptor only occasionally and transiently bind ZAP70-GFP.

We next investigated whether the dynamics of ZAP70-GFP recruitment to a DNA-CAR consisting of all TCR subunits (DNA-CAR_TCR_) was different from DNA-CARζ, which contains just the CD3ζ chain. To generate the DNA-CAR_TCR_, the extracellular SNAP-tag was fused to extracellular domain of the TCR β-subunit; when expressed in the JRT3 cell line (which is null for the β-subunit), the SNAP-β-subunit assembles with the other TCR subunits (α chain, and the ITAM-containing CD3ε, δ, γ, ζ chains) and the full TCR complex is then able to traffic from the ER to the cell surface (Fig. 4A and Fig. S5A-B; Movie S5 and S6)^27^. At a low density of 0.1 per µm^2^, the minority (~21%) of single DNA-CAR_TCR_ receptors bound with 16mer ligand recruited ZAP70 (Fig. 4B, and Movie S7). As was observed with the DNA-CARζ the ZAP70 dwell time at the single ligand-bound DNA-CAR_TCR_ was transient, with a mean dwell time of ~20s (Fig. 4C, examples in Fig. 4D and Fig. S5C). At this low ligand density, we also observed the rare formation of a small cluster consisting of two 16mer ligands (~5% of total events, Fig. 4B). Of these rare events (n = 8), ~63% recruited ZAP70 (Fig. 4B, see example 3 in Fig. S5C), a result that is consistent with a mechanism of small clusters of receptors being more readily phosphorylated than single ligated receptors. When the 16mer ligand density was raised to 1 molecule per µm^2^, clusters of ligand-bound DNA-CAR_TCR_ began to form more readily and these clusters more efficiently recruited ZAP70 (90%) compared with single ligated receptors (20%) (Fig. 4E, examples in Fig. S5D). Furthermore, the ZAP70 dwell time at single ligand-receptor complexes (~20s Fig. 4F) was much shorter compared with clusters; one-third of receptor clusters displayed ZAP70-GFP associations times of >100s (Fig. 4G). These results reveal similar behavior of DNA-CAR_TCR_ and DNA-CARζ; in both instances, small receptor-ligand clusters more efficiently and stably recruiting ZAP70 compared to single ligated receptors.

**Fig. 4.**
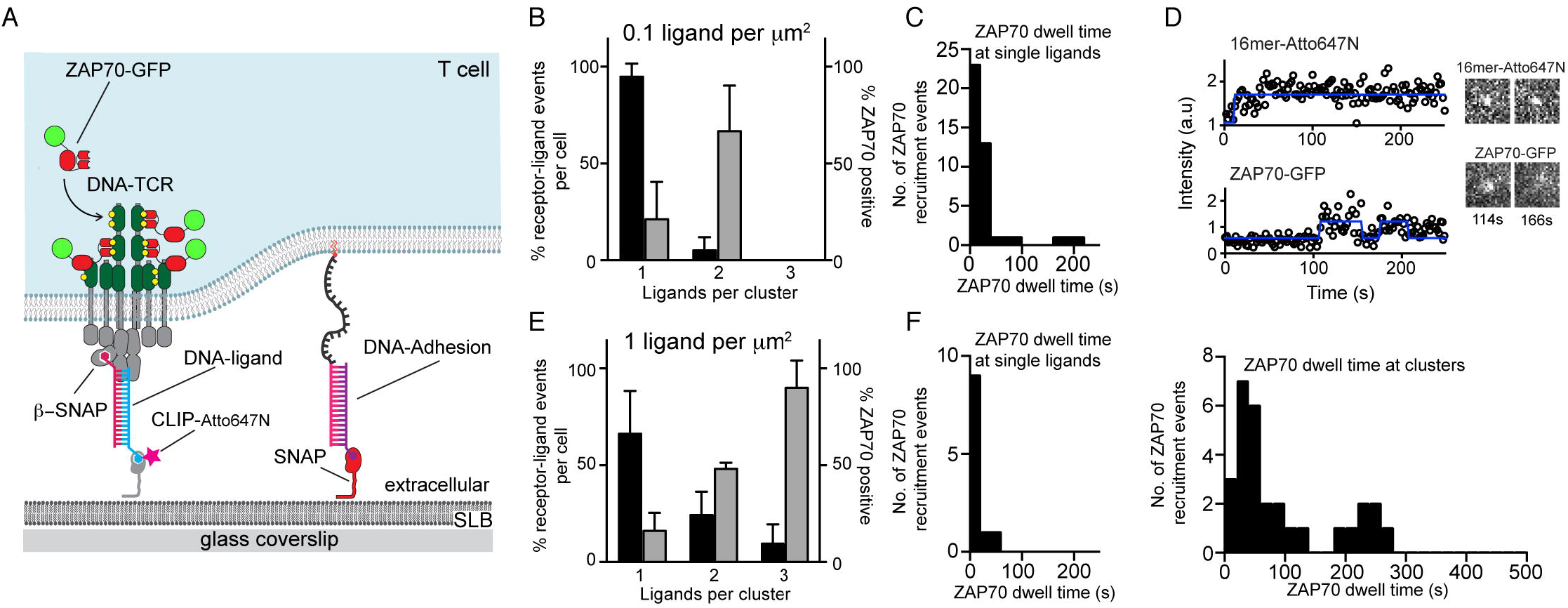
Receptor clusters increase the probability of ZAP70 recruitment to a DNA-based chimeric antigen receptor system consisting of the complete TCR. **A.** Schematic of a DNA-based chimeric antigen receptor system consisting of the complete TCR (DNA-CAR_TCR_). **B.** Quantification of ZAP70-GFP recruitment to DNA-CAR_TCR_ at 0.1 ligand/µm^2^ of 16mer DNA ligand. Bar plot shows the number single bound ligand-receptors and clusters (percent of total, black bars) and ZAP70 recruitment to ligand-receptors (percent of total that recruit ZAP70, grey bars). 21 ± 19% of single bound ligands that persist greater than 30 s recruit ZAP70. Results are mean ± s.d. from n = 8 cells. **C.** Quantification of ZAP70 dwell time at single bound ligands of 16mer DNA ligand (n=42). **D.** A typical example of a transient recruitment of ZAP70-GFP to a single DNA-CAR_TCR_. **E.** Quantification of ZAP70-GFP recruitment to DNA-CAR_TCR_ at 1 ligand/µm^2^ of 16mer DNA ligand. Organization of bar plot same as shown (B). 90 ± 9% of receptor-ligand clusters composed of 3 or greater ligands recruit ZAP70 versus 16 ± 9% of single bound ligands. Results are mean ± s.d. from n = 3 cells. **F.** Distribution of ZAP70 dwell times at single bound receptors (n=11) and receptor-ligand clusters (n=30).

### Comparison of low and high affinity ligands

We next sought to compare the behaviors of a low affinity (13mer DNA strand) and high affinity receptor (16mer) receptor interacting with their cognate ligands at the same density on a supported lipid bilayer (1 molecule per µm^2^). At this ligand density, at which only the higher affinity 16mer elicits a pERK response (Fig. 2B), the two receptors showed considerable differences in their abilities to form clusters. In the case of the 16mer, many (5-20) receptors clusters formed a few minutes of cells after landed on the bilayer (Fig. 5A), and many of these clusters were long-lived (44% persisting for >100 s, Fig. 5B). During the same period of time with the 13mer ligand, single ligand-receptor binding events were observed but cluster formation was very rare (Fig. 5A; Movie S8). A few small clusters of ligated 13mer receptors began to appear on the cell surface after 15 min (Fig. 5A; Movie S9), but most of these clusters disassembled rapidly (mean half-life of ~21 sec; Fig. 5C). As described earlier for the 16mer ligand, ZAP70-GFP recruitment was observed with ~50% for clusters of three or more 13mer ligands and infrequently (~2%) observed with single 13mer ligand-receptor complexes (Fig. S6A, B). Since these 13mer receptors clusters were transient, correspondingly, ZAP70-GFP also dissociated from the membrane (Fig. S6C). At 30-fold-higher 13mer density on the bilayer (30 molecules per µm^2^; where 20% of the cells become pERK-positive (Fig. 2B)), DNA-CARζ clusters now formed within a few minutes of the cell landing on the bilayer (Fig. 5A). These clusters had a similar stability to that seen with the 16mer at 1 molecule per µm^2^ (36% persisting for >100 s, Fig. 5B), and efficiently recruited ZAP70-GFP (Fig. S6C). In summary, the strong and weak ligands formed clusters and stably recruited ZAP70 at different ligand densities; the ligand density required for ZAP70 recruitment also correlated with that required to generate a rapid downstream pERK signaling response (Fig. 2B).

**Fig. 5.**
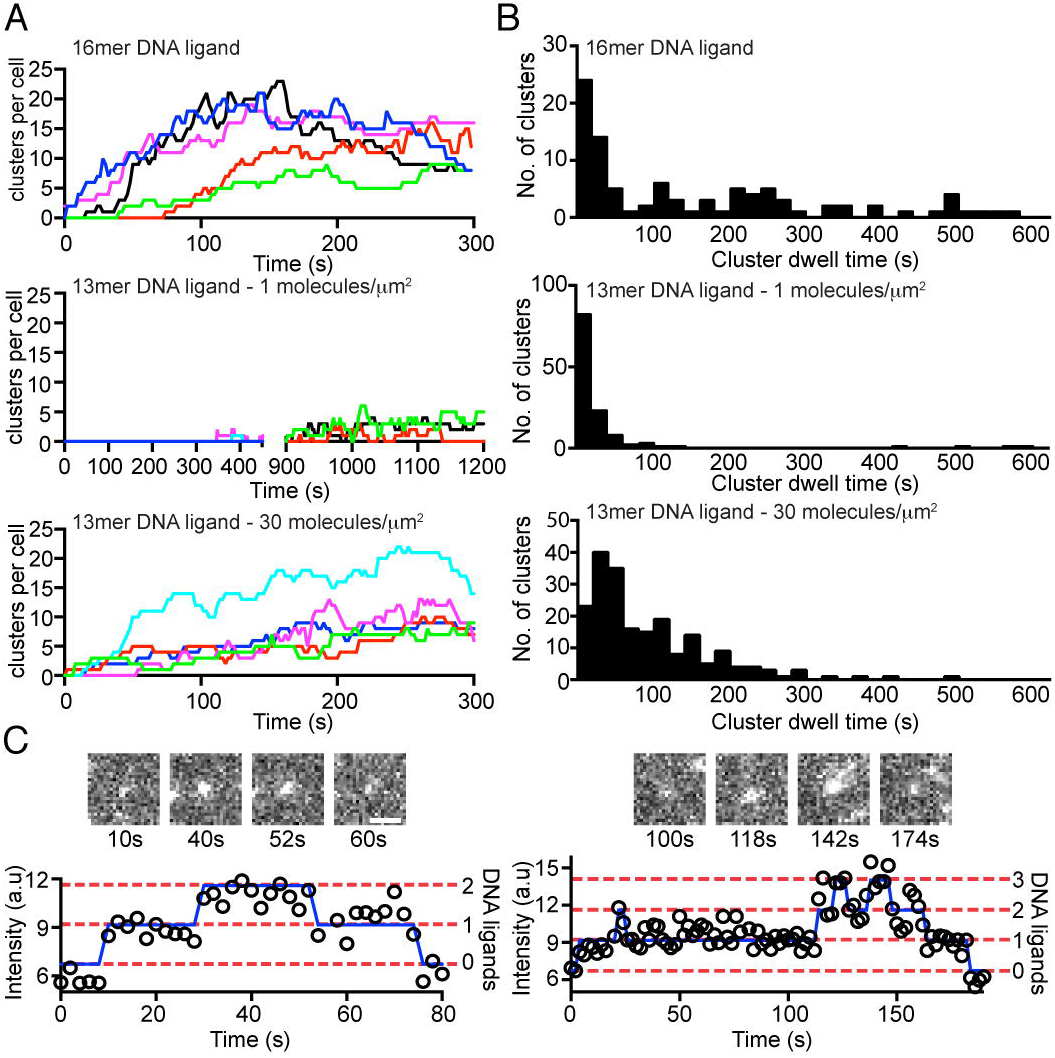
Difference in microcluster formation by low and high affinity ligands with DNA-CARζ. **A.** Formation over time of ligand-receptor clusters (defined as a diffraction limit structure containing 2 or more bound ligands) for individual cells (colors) between at 1 ligand/µm^2^ for the 16mer, 13mer DNA ligand and 13mer DNA ligand at 30 ligand/µm^2^. For 16mer and 13mer plots at 0-15 min, t=0 is defined as the point of image acquisition, often within 1-2 min of application of cells to the SLB. **B.** Distribution of dwell time for ligand-receptor clusters composed of 16mer (n=94 from 6 cells, 35 clusters with dwell times of >100 s and were truncated by the end of image acquisition) and 13mer DNA ligands (n=125 from 6 cells, 6 clusters with dwell times of >100 s were truncated by the end of image acquisition) at 1 ligand/µm^2^ and 13mer DNA ligand at 30 ligand/µm^2^ (n=203 from 5 cells, 26 clusters with dwell times of >100 s were truncated by the end of image acquisition). **C.** TIRF images and intensity time series showing the formation of transient receptor-ligand clusters of 13mer DNA ligand at 1 ligand/µm^2^. The fluorescent time series were analyzed as described in Fig. 3 and Fig. S3.

### Enhanced on-rate of ligand binding adjacent to pre-existing ligand-receptor complexes

We anticipated that receptor-ligand clusters might form through collisions between ligated receptors that are passively diffusing or actively being moved by the actin cytoskeleton in the plane of the plasma membrane. However, such collisions and fusions between single ligated receptors were rarely observed. More commonly, a single DNA-CARζ (Fig. 6A; Movie S4 and S10) or DNA-CAR_TCR_ (Fig. S6D; Movie S11) grew in fluorescence intensity in roughly quantized steps This result is best explained by new ligand binding events occurring near to (within the diffraction limit boundary) pre-existing receptor-ligand complexes. We quantified the rate of new DNA-CARζ-ligand binding events occurring adjacent to a pre-existing receptor-ligand (using an area of a diffraction-limited spot) versus the rest of the plasma membrane (area of the total cell footprint on the SLB observed in the TIRFM field, Fig. 6B). This analysis revealed that the area-normalized on-rate is at least 350-fold higher near a pre-existing ligated receptor or small receptor cluster compared to the rest of the membrane (Fig. 6B). We observed a similar phenomenon of an enhanced ligand binding rate at pre-existing bound DNA-CARζ-ligand complexes in cells treated with latrunculin to depolymerize actin, although the magnitude of the effect was diminished (75-fold enhancement; Fig. S6E-F). After small clusters formed, we observed that they diffused and could sometimes fuse with one another to form larger sized clusters (Fig. 6A; Movie S10). In summary, enhanced ligand on-rate adjacent to pre-existing ligated receptors dominates the formation and growth of receptor clusters. Potential mechanisms that could explain this result are presented in the Discussion.

**Fig. 6.**
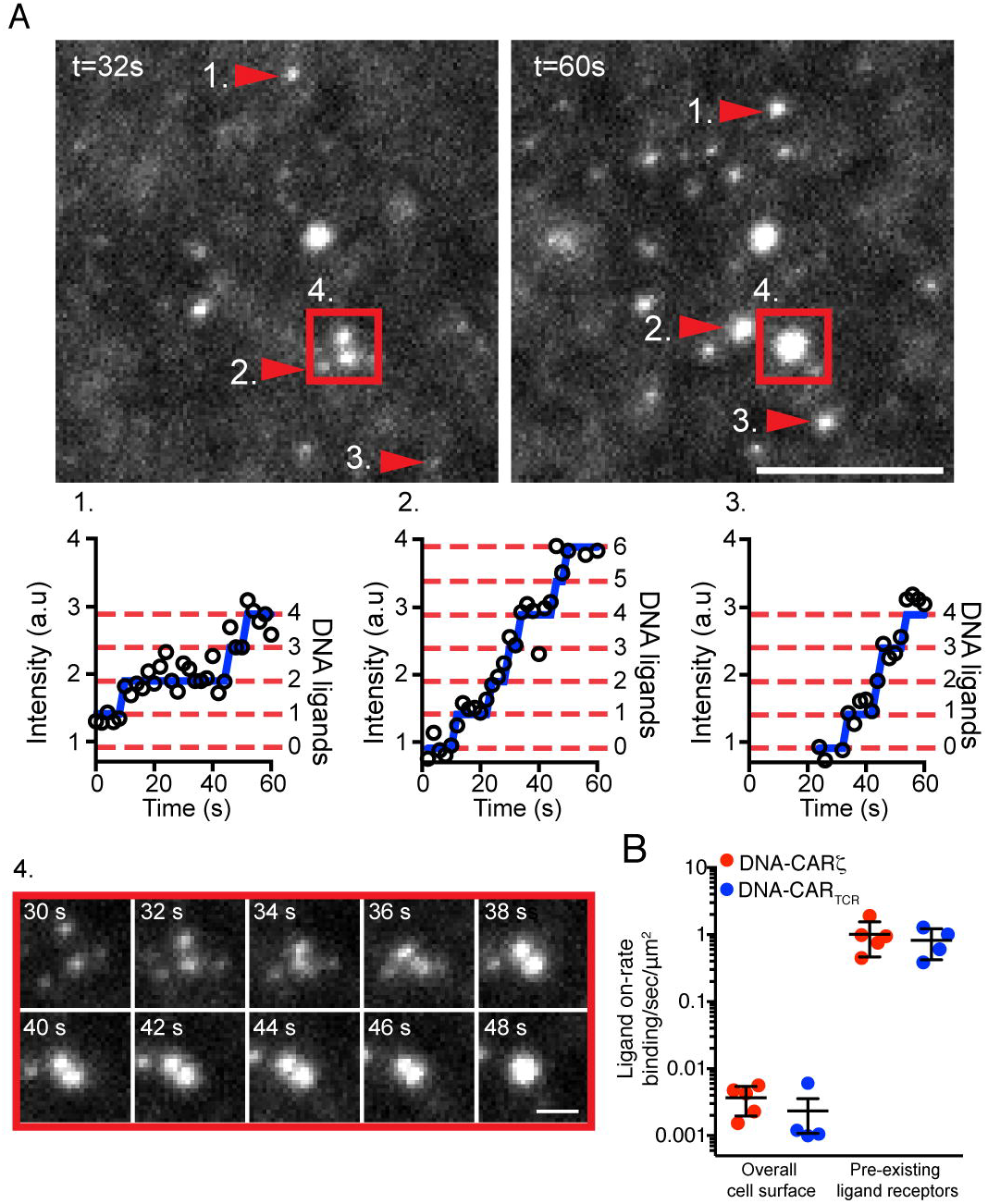
Formation of microclusters from single ligand-receptor binding events. **A.** A TIRF image of receptor-bound 16mer DNA ligand organized into receptor-ligand clusters that grow by adjacent ligand binding events (red arrows numbered 1-3) and by merging and fusion (red box numbered 4). Bar, 2.5 µm. See Movie S10. Clusters grow by sequential addition of newly bound ligand; the blue line overlaid the fluorescent intensity represents detected step changes in fluorescence intensity (see methods and Fig. S3). Time series (red box, numbered 4) below follows two receptor-ligand clusters (bar, 1 µm). **B.** The rate of new ligand binding events near to an existing receptor-bound ligand (quantal increase in intensity in an existing diffraction-limited spot) or outside of these zones (measured by the sudden appearance of a new bound ligand in the contact area between the cell with the supported lipid bilayer) for DNA-CARζ (red) and DNA-CAR_TCR_ (blue). The ligand-receptor on-rate was expressed as the number of events per sec per µ^2^ membrane surface area (using 0.126 µm^2^ for a diffraction limited spot of a pre-existing ligand-receptor pair). Shown is mean ± s.e.m. from 5 and 4 cells for the DNA-CARζ and DNA-CAR_TCR_ respectively.

## Discussion

In summary, our data, for both a chimeric antigen receptor and a complete TCR, show that the binding energy of extracellular DNA hybridization can be transduced across the plasma membrane to trigger intracellular receptor phosphorylation and further downstream signaling. We show that a longer bound-time between a ligand and its receptor and higher ligand densities synergize to promote receptor clustering and that the formation of long-lived receptor clusters substantially increases the probability of receptor phosphorylation and ZAP70-GFP recruitment compared with even long-lived single ligated receptors. These results provide new insights into the mechanism of TCR signaling and the basis of ligand discrimination, as discussed below.

### The Role of Receptor Clustering in TCR Signaling

Using DNA hybridization, we could examine a ligand (16mer) with a much longer predicted off-rate (>7 h) than the strongest agonist pMHCs (~1 min; ^11,21,43^). Surprisingly, despite the long engagement of the 16mer, the majority of single 16mer ligand-receptor pairs that could be tracked for >30 s did not become phosphorylated and recruit ZAP70. In instances where ZAP70 was recruited, its residence time was short (~10-20 s), which is similar to the off-rate measured for ZAP70 from CD3ζ ^17,44^. Thus, we suspect that these transient recruitment events reflect a relatively rare dual phosphorylation event of an ITAM on CD3ζ by Lck and the recruitment of ZAP70, followed by the dissociation of ZAP70 and the rapid dephosphorylation of CD3ζ by CD45 to prevent rebinding of ZAP70. Some models of TCR signaling propose that receptor-ligand dwell time is the primarily determinant of T cell receptor activation ^39,40^. However, our results indicate that a long receptor-ligand engagement per se is insufficient to induce effective downstream signaling through ZAP70 recruitment to the plasma membrane.

Our results provide strong support for the role of clusters in T cell signaling and are consistent with statistical analysis indicating that the small clusters of ligated pMHC-TCR activate downstream calcium signaling ^45^. In contrast to the single ligated receptors, small clusters of 3 ligated receptors are ~90% occupied with ZAP70 and the dwell time of the ZAP70 at the membrane increases substantially. We have observed hundreds of instances of a time-dependent conversion of single ligated receptors into clusters, which then became active in recruiting ZAP70. Interestingly, the dwell of ZAP70 at these small receptor clusters does not follow a simple exponential and a subset of the clusters stably recruit ZAP70 for >2 min (Fig. 3F).

Possible mechanistic explanations for such enhancement in ZAP70 dwell time could be due to dissociation and rapid rebinding to ITAM motifs as a result of high local concentrations in receptor-ligand clusters. Alternatively, clustering might enhance the phosphorylation of ZAP70 at activating tyrosine residues by Lck or other ZAP70 molecules ^46^, a modification that has been shown to enhance ZAP70 affinity for the phosphorylated ITAMs *in vitro* ^17^. We speculate that receptor clusters become more stably phosphorylated than single ligated receptors because they more effectively exclude the transmembrane phosphatase CD45 (Fig. 7A).

**Fig. 7.**
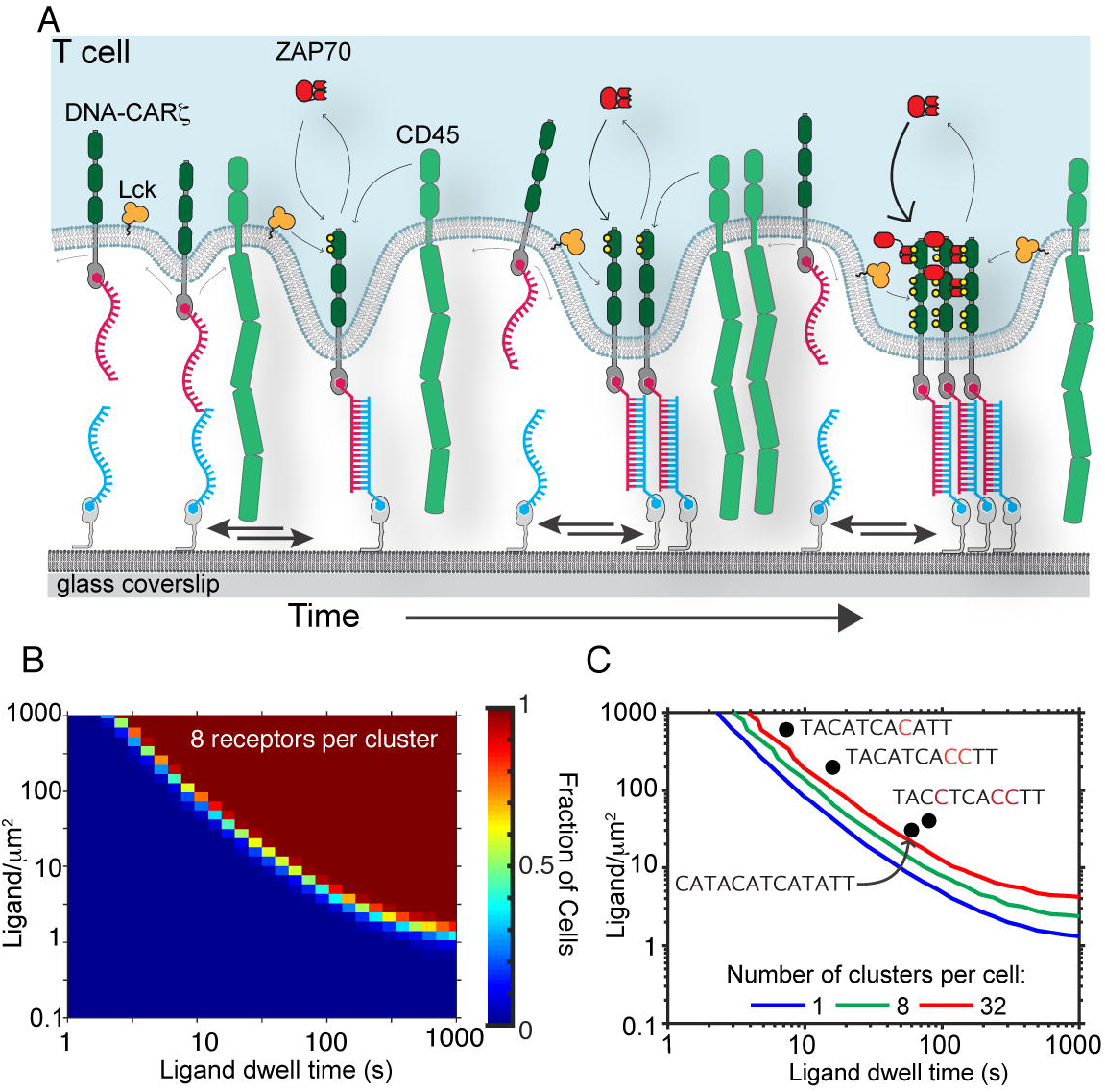
A signaling model for T cell ligand discrimination based on receptor clustering and ZAP70 recruitment. **A.** Model of how a single receptor-ligand binding event nucleates receptor-ligand microclusters which induce receptor phosphorylation. A single receptor-ligand interaction pins the two membranes in close apposition; unbound ligands that diffuse into this region are in closer proximity to and can more readily bind a receptor. Receptors in clusters, which more effectively exclude the phosphatase CD45, become phosphorylated; for lower affinity ligands, receptor clusters and phosphorylation can be reversed by ligand dissociation, providing a mechanism for kinetic proofreading (see text). **B-C.** We constructed a theoretical signalling model, incorporating experimentally measured parameters as described in supplemental Fig. S7, to perform stochastic simulations of cells interacting with a ligand functionalized supported lipid bilayer. These simulations were performed over a range of ligand densities (0.01 to 1000 molecules per µm^2^) and ligand affinities (off-times of 1 to 1000 s) for a fixed time interval of 500 s and repeated for 250 cells at each point in this parameter space. The output of this simulation was time-series data of ligand binding, cluster formation, and ZAP70 recruitment. We analyzed this time series data for appearance of ZAP70 positive receptor-ligand clusters. From this analysis, we generated heat maps showing the fraction of cells where these criteria were observed in a parameter space of ligand density vs. ligand affinity (see Fig. S7K). **B.** Fraction of simulated cells with at least one receptor-ligand clusters containing 8 ZAP70 positive receptors. **C.** Increasing ligand density and affinity results in an increasing number of ligand receptor clusters. The 20% contour of these heat maps was analysed to demarcate regions within this parameter space where the simulation showed >20% of simulated cells forming the indicated number of clusters with a minimum of 8 ZAP70 positive receptors. To compare the predicted emergence of these features in the simulated data with experimental data, we plotted the ligand densities and ligand dwell time at which the indicated DNA ligands elicited a response of 20% phospho-Erk-positive cells (Fig. 2 and S2).

As described in the “kinetic segregation” model^47^, regions of membrane bending created by receptor-ligand interaction exclude the large transmembrane domain of CD45 and thus shift an equilibrium reaction between the receptor kinase (Lck) and the phosphatase (CD45) to favor net receptor phosphorylation. Exclusion of the transmembrane phosphatase CD45 has been observed for receptor clusters composed of many tens or hundreds of molecules ^29,48^. However, if CD45 exclusion underlies the receptor phosphorylation observed in this study, then these results suggest that single ligated receptors are ineffective at preventing CD45 from acting upon phosphorylated TCR, but that clusters even as small as 2-4 receptors can create physical zones that limit access of CD45 phosphatase to phosphorylated ITAMs (Fig. 7A). The expected diameters of the exclusion zones created by these small receptor clusters are well below the diffraction limit of light and most super-resolution light microscopy techniques, but could be examined by electron microscopy in future studies.

The manner in which receptor clusters form also was surprising. Our prior model speculated that single receptor-ligands rapidly diffuse in the plane of the membrane and coalesce into clusters^48^. Instead, this work shows that clusters formed predominantly through an enhancement of the ligand on-rate adjacent to a pre-existing ligated receptor(s). The mechanism behind the dramatic acceleration of the ligand on-rate near to pre-existing ligand-receptors is not established. This observation might arise from heterogeneities in the concentration of the TCR within the membrane, potentially through mechanisms of receptor nano-scale clustering^49^ or dynamic changes in local receptor concentration induced by ligand binding^50^. EM and super-resolution microscopy based studies also have suggested that the TCR is organized into small nano-clusters consisting anywhere between 5-30 receptors^51,52^. However, the nano-scale organization of the unbound TCRs remains controversial with conflicting data ^53,54^, and the mechanism for how nanoclusters of unligated TCRs might assembly and be held together is unknown. Furthermore, we observed enhanced ligand binding with both DNA-CARζ and DNA-CAR_TCR_, which would suggest that, if nano-scale organization exists and is responsible for the enhanced on-rate, it can not be specific to the native TCR.

An alternative explanation for a spatial enhancement in ligand binding evokes the closer physical proximity of the two membranes established by an initial receptor-ligand interaction^3,48^ (Fig. 7A). Theoretical studies have suggested that unbound receptors and ligands on opposite membranes that diffuse into zones of close contact interact more readily, as compared to regions where the two membranes are further apart^55,56^. Computational simulations also have shown that a close membrane contact zone created by an initial receptor-ligand bond facilitates subsequent binding events, resulting in a net cooperative binding effect ^57^. Our experimental results are consistent with the results of this computational study.

### Spatial Organization as a Mechanism of Ligand Discrimination

Like prior studies with different pMHC ligands interacting with TCR ^11,28,58^, we show in this study that T cells can discriminate between DNA ligands with relatively small differences (a few fold) in receptor-bound times (Fig. 2). The molecular basis by which T cells convert such small energy differences in receptor-ligand binding into all-or-none signaling responses and cell activation remains an important unsolved problem. A general model for how this type of ligand discrimination might be generated is called “kinetic proofreading” ^15^. This model proposes that a signaling competent state requires a series of reactions that require the continuous receptor occupancy; if the ligand dissociates, then these reactions are rapidly reversed, terminating signaling and resetting the receptor to its initial inactive state. While this theory is appealing, the series of reversible reactions that lead to a “signaling competent state” are unknown, although many diverse models have been proposed, including receptor conformational changes, dimerization, and phosphorylation reactions^15-20^.

Our results suggest that the spatial organization of receptors provides a mechanism for kinetic proof-reading in ligand discrimination. In line with the kinetic proofreading model, the initial ligand-receptor complex has low signaling output and must undergo a series of time-dependent and reversible steps to form a signaling-competent receptor cluster. Cluster formation is driven by new ligand binding adjacent to a pre-existing ligated receptor, and thus depends upon the ligand density on the membrane (the association rate being concentration dependent) as well as off-rate of pre-existing ligand-receptor complexes. Weaker ligands, as seen with the 13 mer at 1 molecule per µm^2^ (Fig. 5), dissociate faster than new binding events occur, prohibiting the build-up of stable signaling-competent receptor clusters.

To quantitatively assess whether the time-dependent formation of receptor clusters provides a mechanism for ligand discrimination, we constructed a kinetic proofreading mathematical model that incorporates our experimentally determined on- and off-rates of receptor-ligand binding and ZAP70 interaction with the membrane (Fig. S7). In principle, each ligated DNA-CARζ may bind three ZAP70 molecules (one at each ITAM on the CD3ζ cytoplasmic domain); however in this model, we make the simplifying assumption that each receptor is either ZAP70 positive or negative (congruent with our image analysis, Fig. S3E). Using our experimentally derived rate constants, we computed the statistics of the number of ligated receptors in a cluster at the moment of ZAP70 recruitment, the propensity of single receptors or receptor clusters to become ZAP70 positive, and the distribution of cluster lifetimes (Fig. S7G-I). The results of our simulations of the above parameters agreed well with experimental data, thus validating the overall model (Fig. S7).

To understand how the formation of ZAP70 positive clusters could provide thresholds for a discriminatory signalling output, we conducted stochastic simulations ^59^ of the model in a parameter space of ligand density and affinity. We analysed the results of the simulation by plotting the fraction of simulated cells that formed clusters consisting of a few ZAP70 positive receptors (here chosen to be 8; variations of the threshold in Fig. S7J) (Fig. 7B). The results of this simulation reveal a sharp transition (~3 fold changes in ligand density or ligand affinity) between a regime of a low probability of cluster formation (<20% of cells; blue area in Fig. 7B) to a regime of high probability receptor clustering with stable ZAP70 recruitment (>80% of cells) (red/brown area in Fig. 7B, and Fig. S7). We also analyzed simulations that incorporated the very long ZAP70 dwell times as shown in Fig.s 3 and 4. These simulations also revealed similar sharp transitions (Fig. S7L-K) but revealed a greater probability of stable ZAP70 association at clusters of high-affinity ligands at low ligand densities, thus further enhancing ligand discrimination of this kinetic proof-reading model. In summary, our simulations reveal a switch-like response for receptor cluster formation. This behaviour can amplify small differences in ligand affinity or density into dramatically different outputs of ZAP70 recruitment

We next analyzed how well our model correlated with the experimentally observed downstream signalling outputs of phosphoErk (evaluated as 20% phosphoERK activation threshold, Fig. 2B). We found that phospoERK signalling threshold lay inside the region of parameter space in which the signalling model generated multiple clusters that stably recruited ZAP70 (Fig. 7C and Fig. S7J). In the regime for which we measured phosphoERK signalling, the slope of the data and the activation threshold of the model were similar, which reflects a similar degree of ligand discrimination. The model predicts that a 11mer DNA ligand (TACATCATATT), with a ~2 s dwell time (Fig. 2B and D), would generate clusters with stable ZAP70 at a density of ~900-1000 ligand per µm^2^ (Fig. 7L-K), and thus would trigger a phosphoERK response. However, our experimental analysis revealed no discernible signaling for the 11mer ligand (Fig. 2), implying that the degree of ligand discrimination in our mathematical model is lower than that observed experimentally. Thus, kinetic proofreading through receptor clustering provides a partial, but not complete explanation for ligand discrimination, and implies that other kinetic proofreading steps may lie downstream of ZAP70 recruitment to receptor clusters.

### Sensitivity of T Cell Signaling

The bound time of the 13mer DNA receptor-ligand (~24 sec) is comparable to TCR-pMHC complexes that have been extensively studied (few tens of seconds^11^). This DNA ligand also elicited T cell signaling at comparable densities (10-100 molecules per µm^2^) to those used to stimulate native T cells with pMHC on supported lipid bilayers^28,45^. However, in the more physiological context of the APC-T cell conjugate, T cells respond to lower levels of pMHC^14,60^. In these contexts, T cell signaling is likely facilitated by other factors, such as force-induced mechanical changes in the TCR^61^, adhesion molecules that also form signaling complexes (e.g. LFA-1-ICAM1)^62^, and numerous other co-receptors that enhance signaling (e.g. CD4/8, CD28-B7, CD2-CD58)^63^. In future work, these additional components can be added to the DNA-CARζ system to determine if they enhance the sensitivity of signaling and, if so, understand how they affect the assembly and phosphorylation kinetics of receptor clusters as well as influence downstream biochemical events that lead to T cell activation.

## Methods

### Cell Culture

JRT3 Jurkat cells (which fail to express the TCR^27^) and P116 Jurkat cells (which do not express ZAP70^42^) were kindly provided by Art Weiss (UCSF). Both cell lines were grown in RPMI (Invitrogen) with 10% FBS (Invitrogen) supplemented with 2 mM L-glutamine. HEK-293T cells (purchased from the ATCC collection) were grown in DMEM (Invitrogen) supplemented with 2 mM L-glutamine. All cells were determined to be negative for mycoplasma using the MycoAlert detection kit (Lonza).

### Generation of DNA Chimeric Antigen Receptors and ZAP70 constructs

Two versions of the DNA-CARz were constructed: 1) DNA-CARz with a C-terminal cytoplasmic monomeric eGFP (herein termed “GFP”) (pHR-DNA-CARz-GFP), and 2) a non-fluorescent alternate version with a N-terminal HA epitope tag (YPYDVPDYA). The HA tag was inserted to allow cell surface expression levels to be monitored via FACS. Apart from the addition of a HA tag or GFP in these two versions, the receptor was otherwise identical. All primers were purchased from Integrated DNA Technology, and primers longer that 60 nucleotides were ordered as Ultramer oligos. Full details of the construction of the DNA-CARz and DNA-CAR_TCR_ are given below.

### DNA-CARζ-GFP

To construct DNA-CAR-GFP, the human CD3ζ cytoplasmic tails (a.a. 58-164) fused to the transmembrane domain of CD86 (a.a. 236-271) was amplified by polymerase chain reaction. The template used for this PCR was a CD86-CD3ζ chimeric receptor previously described^48^). This produced a DNA fragment with a 5’ 3x gly-gly-ser linker and 3’ BamH1 restriction site. A second PCR amplified SNAPf (from the pSNAPf plasmid, New England Biolabs) with a N-terminal signal peptide (MQSGTHWRVLGLCLLSVGVWGQD) derived from

CD3e. This PCR also introduced a 5’ Mlu1 restriction site and 3’ 3x gly-gly-ser linker (complementary to the first PCR product). A stitch PCR was then set up to produce a final PCR product that was digested with Mlu1 and BamH1 and ligated in frame with mGFP in the second-generation pHR-mGFP lentiviral vector.

### DNA-CARζ-IRESpuro

DNA-CARζ-IRESpuro was constructed using DNA-CARζ-GFP as PCR template. An IDT Ultramer forward primer was designed so that a HA epitope tag was inserted between the signal peptide and the SNAPf open reading frame. This version of the DNA-CARz was digested with Mlu1 and BamH1 and ligated into a pHR lentiviral vector that had a downstream IRES-puromycin resistance cassette (pHR-DNA-CARζ-IRES-puro).

### DNA-CAR_TCR_-IRESpuro

DNA-CAR_TCR_ was constructed using the Jurkat TCRb open reading frame as a template. A primer was designed to PCR amplify a DNA fragment consisting of signal peptide fused SNAPf with a 5’ Mlu1 site and 3’ gly-ser-gly-ser linker (this PCR used DNA-CAR as a template). A second set of primers were designed to PCR amplified the Jurkat TCRb open reading frame (omitting the signal peptide) with 5’ portion of SNAPf ORF and gly-ser-gly-ser linker (complementary to the first PCR product) and a 3’ BamH1 site. A stitch PCR was then set up to produce a final PCR product (signal peptide-SNAPf-TCRβ) that was digested with Mlu1 and BamH1 and ligated into a pHR lentiviral vector that had a downstream IRES-puromycin resistance cassette (pHR-DNA-CAR_TCR_-IRESpuro).

### pHR-TCRα-E2A-SNAPf:TCRβ-P2A-CD3ε-P2A-CD3ζ

To overcome low surface expression of DNA-CAR_TCR_ (see Cell Culture and Reagents, below) a vector was constructed to increase expression of additional Jurkat TCR subunits. We created a multicistronic lentiviral vector where multiple TCR subunits and the SNAPf:TCRβ were separated by 2A ‘ribosome skip’ peptides ^64^. DNA fragments of each of the Jurkat TCR subunits were PCR amplified with primers that added either the E2A or P2A peptide sequences at the 5’ and 3’ terminus. PCR also generated fragments with 15-20 bp overlaps the 5’ and 3’. The pHR lentiviral vector was digested with Mlu1 and Not1. DNA fragments were combined with digested vector and assembled using Gibson Assembly cloning.

### ZAP-GFP

pHR-ZAP70-GFP and pHR-ZAP70-mCherry were as described earlier^48^.

### Lentiviral Production and Generation of Stable Expressing JRT3 Cell Lines

Lentivirus particles were produced in HEK-293T cells by co-transfection of the pHR transfer plasmids with second generation packaging plasmids pMD2.G and psPAX2 (a gift from Didier Trono, Addgene plasmid # 12259 and # 12260). Virus particles were harvested from the supernatant after 48-72 hr, filtered and applied to JRT3 cells overnight. The next day the cells were resuspended in fresh RPMI media and recovered for 3 days. DNA-CAR-GFP expressing JRT3 cells were FACS sorted to generate a stable and homogenous expressing population. JRT3 transduced with pHR-DNA-CARζ-IRES-puro/pHR-DNA-CAR_TCR_-IRESpuro were selected at 4 µg/ml of puromycin (Sigma) and maintained with 2 µg/ml of puromycin.

FACs analysis of DNA-CAR_TCR_ expressing JRT3 cells revealed low surface expression (as compared to wild type E6.1 Jurkats) of the full TCR complex, despite puromycin selection. To increase plasma membrane expression, JRT3 cells were subsequently transduced with pHR-TCRα-E2A-SNAPf:TCRβ-P2A-CD3ε-P2A-CD3ζ to enhance the expression of additional TCR subunits. Second, to selectively sort for cells with high surface expression levels of DNA-CAR_TCR_, JRT3 cells were labeled with SNAP-Surface-647 (NEB) and sorted by FACS.

This resulted in a population with a plasma membrane expression level of DNA-CAR_TCR_ that was comparable to wild type Jurkats TCR levels (as compared by FACS analysis).

### Imaging Chambers and Supported Lipid Bilayers

Phospholipid mixtures consisting of 97.5% mol 1-palmitoyl-2-oleoyl-*sn*-glycero-3-phosphocholine (POPC), 2% mol 1,2-dioleoyl-sn-glycero-3-[(N-(5-amino-1-carboxypentyl) iminodiacetic acid) succinyl] (nickel salt) (Ni^2+−^NTA-DOGS) and 0.5% mol 1,2-dioleoyl-sn-glycero-3-phosphoethanolamine-N-[methoxy(polyethylene glycol)-5000] (PE-PEG5000) were mixed in glass vials and dried down under argon. All lipids used were purchased from Avanti Polar Lipids. Dried lipids were placed under vacuum for 2 hr to remove trace chloroform and resuspended in PBS. Small unilamellar vesicles were produced by several freeze-thaw cycles. Once the suspension had cleared, the lipids were spun in a bench top ultracentrifuge at 65,000xg for 45 min and kept at 4° C for up to 5 days.

Supported lipid bilayers were formed in 96-well glass bottom plates (Matrical), which were cleaned by extensive rinsing in isopropanol following by water. Plates were then cleaned for 15 min with a 1% Hellmanex solution heated to 50°C followed by extensive washing with pure water. 96 well plates were dried with nitrogen and sealed until needed. To prepare SLB, individual wells were cut out and base etched for 5 min with 5 M KOH and then washed with water and finally PBS. SUVs suspension were then deposited in each well and allowed to form for 1 hr. We found that SUVs suspension containing 0.5% mol PE-PEG5000 formed best at 37°C. After 1 hr, wells were washed extensively with PBS. SLBs were incubated for 15 min with HEPES buffered saline (HBS: 20 mM HEPES, 135 mM NaCl, 4 mM KCl, 10 mM glucose, 1 mM CaCl_2_, 0.5 mM MgCl_2_) containing 1% BSA to block the surface and minimize non-specific protein adsorption. After blocking, the SLB were functionalized by incubation for 1 hr with his-tagged proteins. The labeling solution was then washed out and each well was completely filled with HBS with 1% BSA. Total well volume was 625 µl (manufacturers specifications), and 525 µl was removed leaving 100 µl of HBS 1% BSA in each well.

### Protein Expression, Purification and Labeling

SNAPf and CLIPf open reading frames were cloned into a pET28a vector containing a N-terminal 10X His tag. A C-terminal ybbr13 tag (DSLEFIASKLA)^65^ was added by PCR. Proteins were expressed in BL21-DE3 E. coli and purified by Ni-NTA resin followed by gel filtration. Ybbr13 peptide labeling was performed using CoA-Atto647N as described^65^. The degree of labeled was calculated with a spectrophotometer by comparing 280nm and 640nm absorbance (usually 85-95% labeling efficiency was achieved).

### Synthesis of Benzylguanine-Conjugated DNA Oligonucleotides

All receptor/ligand/adhesion oligonucleotides were ordered from IDT with a 3’/5’ terminal amine. Conjugation to benzyl-guanine or benzyl-cytosine was performed as described ^66^. 10x His tagged SNAP and CLIP were labeled with benzlyguanine/benzylcytosine DNA on the same day SLBs were prepared. SNAP/CLIP were labeled at a concentration of 5 µM with a 3-fold excess of BG/BC-DNA in 20 mM HEPES (pH 7.4) 200 mM NaCl and 1 mM TCEP. DNA-SNAP/CLIP linkage was monitored by mobility shift assays using SDS-PAGE. Maximal labeling was achieved after 40 min at room temperature (~90% labeling efficiency).

### DNA Ligand and Receptor Sequence Design

We selected a 16-nucleotide DNA strand that had no discernable secondary structure (as measured using nupack.org^35^, accessed between 12/2012 and 06/2013) and have been previously characterized^67^. The receptor/ligand 16mer DNA strand had the following sequences: ligand (5′-CCACATACATCATATT-3′) and receptor (5’-AATATGATGTATGTGG-3’). All receptor/ligand DNA strands were ordered from Integrated DNA Technology with 3’ amine functional group. Truncations of the initial 16mer DNA ligand sequence were generated from the 5’ end. To maintain the overall length of the DNA ligand as presented on the SLB, a poly-thymine spacer was added back at the 3’ end. The truncation ligands had the following sequence: 13mer ligand (5’-CATACATCATATTTTT-3’), 12mer ligand (5’-ATACATCATATTTTTT-3’), and 11mer ligand (5’-TACATCATATTTTTT-3’). For experiments using shorter complementary DNA ligands the same 16mer DNA receptor sequence was used (5’-AATATGATGTATGTGG-3’). Mutant versions of the 11mer were generated by sequential addition of C/G base pairs. Mutant 11mer strands were analyzed by nupack.org to minimize secondary structure. The following mutant 11-nucleotide DNA receptor ligand pairs were used (mutations underlined): DNA receptor (5’-AATGTGATGTATTTTT-3’), DNA ligand (5’-TACATCACATTTTTTT-3’), DNA receptor (5’-AAGGTGATGTATTTTT-3’), DNA ligand (5’-TACCTCACCTTTTTTT-3’) and DNA receptor (5’-AAGGTGAGGTATTTTT-3’). The triggerable DNA signaling system used the same 16mer DNA receptor (5’-CCACATACATCATATT-3). The SLB was functionalized with 18mer non-complementary oligonucleotide (5’-CCCTCATTCAATACCTCA-3’). Receptor and ligand were brought together by the addition of an oligo with complementary regions to both receptor and ligand (5’-AATATGATGTATGTGGttCCTAGGGTATTGAATGAGGG-3’). The addition of trigger strand results in a 34 base pair overlap. While this trigger strand system was useful for investigating kinetics by synchronizing the timing of ligand-receptor interaction, the complexity of this three-component binding interaction system^68^ made it difficult to use for generating ligand dose responses.

### DNA-Based Adhesion System

The 100mer adhesion strand used in this study consisted of a 3’ 20mer complementary region (5’-ACTGACTGACTGACTGACTG-3’) attached through a 80mer poly-dT linker to a lipid anchor (1,2-O-Dihexadecyl-sn-glycerol) via a phosphodiester linkage at the 5’ end. Dialkylglycerol phosphoramidites were synthesized as previously described^32,69^. The complementary sequence (5’-CAGTCAGTCAGTCAGTCAGT-3’) was ordered from Integrated DNA Technology with a 5’ amine and labeled with BG-NHS as previously described above. This strand was then conjugated to His10-SNAPf to label SLBs. Cells were labeled with the DNA adhesion lipid molecule for 3 min at room temperature at a labeling concentration of 5 mM (stock concentration of 250 mM).

### BG-DNA Labeling of JRT3 Cells Expressing DNA-CARz

JRT3 cells expressing DNA-CARz/DNA-CAR_TCR_ were spun down, re-suspended in HBS and incubated with 5 µM of BG-DNA receptor for 30 min at room temperature. Cells were conjugated with BG-DNA at an approximate density of 2 × 10^7^ cells/ml. During conjugation cells were maintained in suspension by gently agitation. DNA adhesion lipid was added during the final 3 min of labeling. Cells were washed twice in HBS before being used.

### CD69 Expression

To assay CD69 expression by FACS, supported lipid bilayer were set up on 7 µm silica beads (Bangs Laboratories). Silica beads were counted using a hemocytometer mixed with 2.5 × 10^5^ JRT3 cells expression DNA-CARz-GFP in 96 well plates in a 3:1 ratio of bead to cells. Signaling was initiated by the addition of DNA trigger strand. A portion of cells were also plated onto poly-L-lysine containing coverslips and analyzed by spinning disk confocal to inspect SLB quality and confirm cellular activation via re-localization of DNA-CAR-GFP to the bead-cell interface after addition of the trigger DNA strand (Fig. S1C). 4 hr after activation cells were pelleted and re-suspended in PBS with 2% (v/v) fetal bovine serum and 0.1% (w/v) NaN_3_. Cells were labeled with mouse anti-CD69 conjugated to Alexa647 (FN50, Thermo Fisher Scientific, 10 mg/ml) for 1 hr on ice. Cells were washed twice and then fixed. Cells were then run on a LSRII (Becton Dickinson) (10,000 gated cells analyzed).

### Calcium Imaging and Analysis

Calcium signaling assay was performed on JRT3 cells pre-incubated with 10 µM Fura-2 (Invitrogen) for 30 min. Ratiometric Fura-2 imaging (340 nm/380 nm excitation) was performed on a microscope (Nikon TE2000U) equipped with wavelength switcher (Sutter Instrument Co. Sutter Lambda XL lamp) and Fura-2 excitation and emission filters. Images were projected on to Photometric CoolSNAP HQ2 CCD camera using an S Fluor 40X 1.3 NA oil objective. JRT3 cells expressing DNA-CARz-GFP were pipetted onto supported lipid bilayer incubated for 10 min to allow cells to settle and adhere to the SLB using the DNA adhesion system described. Cells were imaged for 1-2 min in a quiescent state before the addition of trigger strand to initiate signaling. Image analysis was performed in FIJI^70^ by manually segmenting the cell outline and measuring the mean 340 nm/380 nm excitation ratio in the cell volume.

### PhosphoERK Data Acquisition and Analysis

Titrations of ligand density on SLBs were set up using 96 well plates. For each phosphoERK assay, all SLB ligand densities were set up in triplicate. Ligand density was determined by maintaining identical labeling protein concentrations and time, but changing the portion of DNA-ligand labeled His10-CLIPf-Atto647N. Before application of cells, SLBs were analyzed by TIRFM to check formation, mobility and uniformity. Short time series were collected at low ligand densities (e.g. ≥1 molecule per µ^2^) to calculate ligand densities on the SLB based upon direct single molecule counting. Wells containing only DNA adhesion strands served as unstimulated controls or used for phorbol 12-myristate 13-acetate stimulation.

On the day of an experiment JRT3 cells expressing DNA-CARz-GFP were transferred to serum free media for several hr before being functionalized with BG-DNA receptor. After DNA functionalization cells were re-suspended in HBS at a final concentration of 2.5 × 10^5^ cells per ml. 100 µl of cells (corresponding to 2.5 × 10^4^ cells per well) were then applied to 96 plates wells using a multichannel pipette (total well volume after addition of cell was 200 µl). Cells were then incubated at 37°C for 15 min before the addition of 200 µl of 2x fixative (7% (v/w) PFA with 1% (v/w) Triton X). Cells were fixed for 20 min at room temperature. Cells were then washed with PBS containing 60 mM glycine to quench PFA. Cells were then blocked in PBS 10% (w/v) BSA for 1 hr before addition of primary antibody. Cells were labeled over night with anti-phosphoERK (rabbit polyclonal, Cell Signalling Technology #9101, used at 1:500). The next day cells were washed 5X in PBS 10% (w/v) BSA, and labeled with goat anti-rabbit conjugated to Alexa555 (Invitrogen, used at 1:1000) for 1 hr. Finally cells were washed 5X in PBS. In the penultimate PBS wash, cells were labeled for 10 min with DAPI at a labeling concentration of 300 nM.

96 well plates were imaged on an inverted microscope (Nikon TiE, Tokyo, Japan) equipped with Lumencor Spectra-X illumination. Fluorescent images were acquired with Nikon plan apo 20X 0.75 NA air objective lens and projected on an Andor Zyla 5.2 camera with 2x2 binning (pixel size 425nm) and a 1.5x magnification lens. The fluorescent emission was collected through filters for EGFP (525 ± 30nm), Alexa 555 (607 ± 36nm) and DAPI (440 ± 40). Image acquisition was performed using MicroManager software^42,71^. Each well was imaged using the Create Grid plugin in the MicroManager multidimensional acquisition GUI. The Create Grid plugin was used to automate the acquisition of the entire well. A dark image was subtracted from each image during acquisition using the Multi-channel shading MicroManager plugin.

Image analysis was performed using Cell Profiler^38^ and FIJI. Unsuitable images that had focus defects or fluorescent debris were discarded from the image series from each well. The Alexa555 channel, corresponding to the phosphoERK staining, was processed in FIJI using the roll ball background subtraction (ball size 100 pixels) to create a background image. Background images from multiple fields of views were averaged to create an image of the illumination function of the microscope. Each Alexa555 image was then divided by this illumination image using the Cell Profiler plugin ‘Correct illumination apply’ to correct for illumination defects.

To score phosphoErk positive cells, selected phosphoERK and DAPI images from individual wells were processed in batch using a custom Cell Profiler pipeline. The DAPI channel was segmented to identify cell nuclei. The segmented nuclei were used to seed a second segmentation of the phosphoERK stained channel (Fig. S2A). Thresholding parameters for the phosphoERK channel were set using images of PMA stimulated cells, unstimulated cells, and cells labeled with secondary only (for background fluorescence). Segmented nuclei were then related to the segmented phosphoERK objects to score nuclei as phosphoERK positive or negative (e.g. nuclei associate with or without a phosphoERK object). Due to the presence of small phosphoERK positive foci in a portion of cells (found in a fraction of cells even without ligand stimulation), we stipulated that phosphoERK segmented object had to have a minimum size (a minimum diameter of 20 pixels, Fig. S2A). This selected for a phosphoERK staining that had an equivalent size and morphology to the DAPI stain, and was equivalent to the phosphoERK staining morphology of PMA-stimulated cells. In general, between 2500-5000 cells were analyzed per well. Positive and negative phosphoERK nuclei were summed across images from the same well.

### Imaging and analysis of single ligand-receptor dwell time

Single molecule measurements of receptor ligand dwell time were performed on an inverted microscope (Nikon TiE,Tokyo, Japan) equipped with a spinning disk confocal and TIRF combined system (Spectral Diskovery, Ontario, Canada). Two colour simultaneous TIRF laser illumination with 488 and 638 nm was provided by directly modulated lasers combined into a two fiber output (Spectral ILE, Ontario, Canada). Following the general methodology of O’Donoghue et al.^21^, single molecule TIRFM measurements were imaged in streaming mode with a 500 ms exposure time to detect the bound fraction of ligand on the supported lipid bilayer. By using a 500 ms exposure, the bound ligands were detected as discrete spots of fluorescence intensity due to relatively slow diffusion of receptor-bound ligands; unbound ligands on the supported lipid diffused much faster and created a background blurred image on the camera detector (see Fig. S2C-D). Fluorescent emissions of GFP (receptor) and Atto647N (ligand) were split using a 650 nm long pass dichroic onto two Andor iXon Ultra EMCCDs (Belfast, Ireland). Illumination was controlled using digital control boards (Arduino Uno, Turin, Italy) and triggers from the cameras. Image acquisition was performed using MicroManager software^71^. A standard constant temperature of 37°C was maintained using an OKO Labs stage top incubator.

Single molecule diffraction limited spots in the far-red channel were detected and tracked using the FIJI plugin “Trackmate”. Ligand dwell times, as computed from the track duration, were fit to a single exponential decay in Prism Graphpad software to calculate τ_obs_, the mean observed dwell time. The dwell times for the four 11mer mutant oligonucleotides with increasing G/C content were tested on the one experimental day. Per each experiment, single molecule measurements were made from between 8-12 cells per DNA ligand. Bleach rates were determined by absorbing His10-CLIPf-ybbr13-Atto647N to clean glass imaged using identical illumination and acquisition conditions, and were obtained on the same day as ligand dwell time measurement. Bleaching data for single molecules was processed and analyzed in an identical manner to ligand dwell time data to determine the rate of bleaching (τ_bl_)(see also Fig. S2). τ_obs_ is a combination of the rate of dissociation and photobleaching, and can be corrected to obtain tcorr using the following formula:

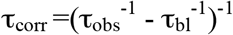

### Imaging and Analysis of Single Ligand-Receptor Interactions and ZAP70 Recruitment

Imaging of ZAP70-GFP recruitment was performed on an inverted microscope (Nikon TiE, Tokyo, Japan) equipped with NIKON fiber launch TIRF illuminator. Illumination was controlled with an Agilent Laser combiner using the 488 and 640 nm laser lines at approximately 0.1 and 0.05 mW laser power respectively. Fluorescence emission was collected through filters for GFP (525 ± 25 nm) and Atto647N (700 ± 75 nm). All images were collected using a Nikon Plan Apo 100x 1.4 NA oil-immersion objective that projected onto a Photometrics Evolve EM-CCD camera with a calculated pixel size of 103 nm. A constant temperature of 37°C was maintained using a Tokai Hit stage top incubator.

JRT3 cells expressing DNA-CARz or DNA-CAR_TCR_ (under puromycin selection) and ZAP70-GFP were pipetted onto supported lipids bilayers functionalized with His10-CLIPf-Atto647N conjugated to DNA ligand. JRT3 cells and SLBs were sequentially illuminated for 500 ms with 488 nm and 640 nm laser lines. Diffraction-limited punctae of Atto647N representing bound DNA ligands were detected and tracked using Trackmate FIJI plugin as described above. A hidden Markov Model (HMM) analysis was then used to identify the number of fluorescent ligands in each frame from the fluorescent intensity of a tracked Atto647N ligand cluster. The same analysis was also used to detect the moment ZAP70 was recruitment to a DNA ligand microcluster. The HMM analysis implemented in this study was the statistical maximum evidence approach described previously by Bronson et al. (see Fig. S3 and below).

To analyze ligand on rate, we segmented the cells-SLB interface using the ZAP70-GFP fluorescence by applying a threshold using FIJI. We calculated the cell-SLB interface surface area from the threshold image and using the Analyse Particle plugin, and calculated the median surface area during the initial 3 min of the cell landing on the SLB. We used this to calculate a ligand binding on-rate based on cell-SLB interface area and the number of de novo single molecule events detected in this 3 min window. We calculated the ligand on-rate in clusters by scoring new binding events that occurred after the initial single molecule binding event that seeded that receptor-ligand cluster. The time interval was calculated from the molecule binding event that preceded the subsequent binding event (e.g the time interval between the second and third binding event). Micro-cluster area was estimated as a diffraction limit spot (calculated using a spot with 0.2 µm radius – radius^2^ × π). We only analyzed events where clear quantal steps were detected. In most examples this meant we could reliably analyze the second and third ligand binding events, but in some case we could analyze up to 5 binding event at an individual micro-cluster.

## Quantification and Statistical Analysis

### PhosphoERK Data Analysis

For outcome assessment of phosphoERK activation (Fig. 2 and Fig. S2A-B) experiments were automatically analyzed by Cell Profiler to minimize potential human bias. In Fig. 2B-C each data point represents the mean from one experiment, where each ligand density was measured in triplicate (Mean ± s.d. (n = 3)).

### Hidden Markov Model analysis of receptor-ligand cluster assembly and ZAP70 recruitment

The number of fluorescent ligands in a cluster is well described by a Markov process - that is, a stochastic process of ligand addition (i.e. the binding rate) and rates of ligand 'removal' (i.e. the combination of the unbinding and the bleaching rate). Therefore, we applied Hidden Markov Model methods to analyze the Atto647N channel data (as described in Fig. S3). We implemented this analysis in Matlab by using the software vbFRET (available at http://vbfret.sourceforge.net/ accessed on September 2015). First the intensity time-series of each tracked cluster was extracted from the coordinates generated by TrackMate. We also extracted the intensity values from the five frames that preceded the appearance of the object (to accurately sample background (i.e. no ligand) intensity values). The fluorescent intensity for each tracked microcluster from a cell was then concatenated to create an ensemble time series which was analyzed by the vbFRET software package, which identified the rates of ligand binding and unbinding (or bleaching) and the fluorescence distributions for cluster composed of n=1,2,3… ligands. Finally vbFRET reconstructed the time-series of ligand number for each cluster using the Viterbi algorithm. We manually verified the reconstruction for every cluster in each cell, manually correcting for overfitting (i.e. the assigning of multiple Markov Model states to what is manually identifiable as a single fluorescent intensity state). To assay for the robustness of this analysis to experimental noise we used inverse transform sampling to re-noise a time-series of ligand number from an analyzed experimental dataset. This procedure randomly samples the fluorescent intensity distribution identified by vbFRET from the experimental data, and ensures the reconstructed data accurately reflects the experimental noise (Fig. S3D).

The same analysis protocol was implemented in the ZAP70 channel with minor differences (Fig. S3E). The tracking output of bound ligand coordinates was used to pull out the equivalent fluorescent intensity in the ZAP70-mEGFP channel using a custom written Matlab script. To aid analysis of the ZAP70-GFP signal, we analyzed intensity values extracted from the ZAP70-GFP channel after a rolling ball background subtraction (performed in FIJI with ball size of 3 pixel) in parallel to the raw intensity values. HMM analysis of the ZAP70 data served as a guide for a subsequent careful manual analysis of the data. Manual verification was used to confirm positive recruitment as a puncta of ZAP70-GFP that co-localized and co-migrated with an object in the ligand channel (Fig. S3F).

### Construction of a stochastic signaling model

## Model Construction and Validation

The goal of our theoretical model was to quantitatively assess the degree of ligand discrimination provided by the experimentally observed mechanisms. This model considered a T cell interacting with a ligand functionalised supported lipid bilayer (Fig. S7A) and the transitions that occur between single bound ligands and receptor-ligand clusters (Fig. 7B,C) over a defined passage of time (fixed at 500 s). We fixed the parameters used in our theoretical model directly from the experimental measurements. These parameters were the following: k_b_ is the rate at which new ligand bound to a site consisting of a existing contact site (also the rate at which single receptor bounds ligands converted into clusters). This rate was taken to be proportional to the ligand concentration on the supported lipid bilayer. The constant of proportionality between k_b_ and the ligand concentration [L] was computed by extracting the value of k_b_ from the 16mer data at 1 molecule per mm^2^ data (Fig. S7D). k_u_ is the unbinding rate of ligands, the inverse of which is the average dwell time τ_u_, and was obtained from the single-molecule dwell-time data (Fig. 2 and Fig. S2). k_0_ is the rate at which receptors on the T cell surface bind to ligands on the SLB, forming a *contact site* between the cell and the supported lipid bilayer, referred to as ‘*de novo* binding' (Fig. S7E). 
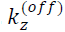
 is the unbinding rate of ZAP70 estimated from the experimental data of ZAP70 dwell times at single ligands (Fig. 3C). 
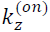 is the on rate of ZAP70 binding and was calculated by measuring the average time between contact site formation and initial ZAP70 recruitment (<T>, Fig. S7F, left), and then inferring the corresponding value of 
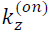 (Fig. S7F, right).

In principle, each DNA-CAR may bind three ZAP70 molecules (one at each ITAM on the CD3z cytoplasmic domain); however in this model, we make the simplifying assumption that each receptor is either ZAP70 positive or negative (congruent with our image analysis, Fig. S3E). Kinetic proofreading arises from this model because ligand unbinding from a ZAP70-positive receptor results in the loss of both a ligand and a ZAP70-positive receptor from a contact site ( (*n, m*) → (*n* − 1, *m* − 1), see state space in Fig. S7B). This loss may only be reversed by two steps ( (*n* − 1, *m* − 1) → (*n, m* − 1) and (*n, m* − 1) → *(n, m)*, i.e. ligand binding followed by ZAP70 recruitment), thereby inducing a temporal delay and energetic cost that is the defining feature of kinetic proofreading ^72,73^. If all ligands unbind from a contact site, then that contact site ceases to exist (i.e. the cluster disassembles). With all parameters fixed, we assess the validity of the model by quantitative comparisons of its predictions with experimental data. Here, we present three such comparisons:

- **Number of 16mer Ligands at Initial ZAP70 Binding:** We plot the prediction of the theory (analytically calculated from the model) against the experimental data (Fig. S7G) and observe an excellent agreement. Note that the data considered here only includes the 16mer tracks that *do* recruit ZAP70.
- **Propensity of 16mer Ligand Clusters to Stay ZAP70-free:** To analyse the ability of the model to understand all of the data, including those tracks that do not recruit ZAP70, we do the following: from the ligand channel of each contact site in the 16mer dataset we compute the probability that the track *does not* recruit ZAP70 (using the assumptions and rates of the model) using Poisson statistics:

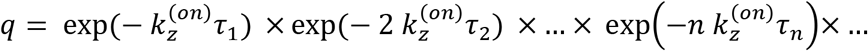

where n is the length of time for which the contact site has exactly n ligated receptors. We then bin contact sites by their q values (Fig. S7H) we have used ten bins (0 → 0.1, 0.1 → 0.2 etc., plotted on the x-axis). For the tracks in each bin, we then ask what fraction actually remains ZAP70-free - plotted on the y-axis. We expect from the model that the data points would fall on the x = y diagonal (black line). We indeed find that the data-points lie close to the diagonal, tracking it very well, but lie consistently above it. This suggests that there is slightly less ZAP70 than expected, which could be due to the difficulty in detecting ZAP70 against the fluctuating fluorescent background, or the recruitment of non-fluorescent ZAP70.
- **13mer Cluster ‘Stability’:** To validate our model in the context of a rapidly unbinding ligand, we look at the ‘lifetime’ of a cluster of 13mers. To be more precise, we consider the amount of time for which the number of ligands at a contact site is n > 1. As the rate of ligand unbinding (
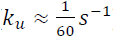) is comparable to the rate of bleaching (
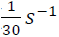), we considered the bleaching-renormalised ligand ‘unbinding’ rate: 1/60 + 1/30 = 1/20 *s*^−1^. The results (obtained from stochastic simulation of the model, Fig. S7I) show an excellent agreement with experimental data.

### Ligand Discrimination in the Model

We performed stochastic simulations^59^ of cells interacting with an SLB of ligand concentration [L] and off-time *τ_u_* = 1/*k_u_*. At each point in parameter space, we simulated N=250 cells for 500 s (chosen to match the experimental timescales); the time-series of (*n, m*) for each receptor-ligand contact site in each cell was then analysed for the following potential *activation thresholds:*

- A minimum cluster size (e.g. number of ligated receptors within a cluster): *n*_*_= 4, 8 or 16.
- A minimum number of ZAP70-positive receptors within a cluster: *m*_*_= 4, 8 or 16.
- A minimum length of time for which a bound ligand or cluster has m>0 ZAP70-positive receptors: t[m]= 100 seconds.

The rationale was to analyse how the emergence of these features correlated with the experimentally observed downstream signalling outputs of phosphoErk (evaluated as 20% phosphoERK activation threshold, Fig. 2B-C). If a cell contained at least one contact site that satisfied the particular activation threshold within the simulation time of 500 s, the cell was scored as having satisfied that threshold criterion. We could then construct *heat maps*, (as in Fig. 7B), that assigned to each point in parameter space the fraction of cells that satisfied the threshold criterion, the 20% contour of which was used to demarcate boundaries of activation in parameter space (Fig. 7C and Fig. S7J-L). The results demonstrate how these activation thresholds can tune the sensitivity and specificity of ligand discrimination. We found that phospoERK signalling thresholds (Fig. 2B-C) mapped to regions in the parameter space were the signalling model found larger clusters that stably recruited ZAP70 (Fig. 7C and Fig. S7J). However by these criteria our model predicts that a 11mer DNA ligand (TACATCATATT) with a ~2 s dwell time and no discernable signalling activity (Fig. 2B and D) would have an affinity sufficient to activate phosphoERK at high ligand concentrations (~900-1000 ligand per mm^2^, Fig. S7J-K). This suggests that the degree of ligand discrimination of this model is lower than experimentally observed, and suggest of other possible mechanisms involved in discrimination.

We went on to analyse the effect of incorporating a ‘cooperative switch’ in ZAP-70 stability, defined as a change in the off-rate of ZAP70 from 
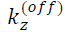 to 
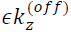 (∊ < 1, see schematic in Fig. S7L) when the number of ligated receptors at a contact site crosses some threshold (here chosen as n > 2). We find (Fig. S7L) that increasing the stability of ZAP70 results in an enhanced sensitivity for high affinity ligands (i.e. shifting the activation boundary for the *m*_*_ or *τ*[*m*] thresholds to lower [L] specifically for ligands with larger off-times *τ_u_*).

### Stochastic Simulation Method

Simulations were performed with the Gillespie stochastic simulation algorithm, implemented in custom C code, in two steps:

- *de novo* binding times i were generating using the rate *k_0_*, between t=0 and t=500s.
- For each of these, a Gillespie simulation was run on the state space in Fig. S_(b), starting in the state (1,0) at t=i

Each contact site was simulated until one of three events occurred: (a) all ligated receptors unbind (n=0), or (b) total simulation time t=500 s was reached.

## Acknowledgments

We thank J. James (LMB and Cambridge University) for initial help and guidance with this project. We thank Sam Lord and Dyche Mullins for use of their microscope. We thank N. Stuurman for assistance with microscopy, C. Carbone for advice and assistance with experimental work, Noel Gee for supplying lipid-modified DNA, and E. Hui and X. Su for comments on the manuscript. Some Image acquisition was performed at the Nikon Imaging Center at the University of California San Francisco, and at the NCBS (Bangalore). R.D.V. is a Howard Hughes Medical Institute investigator. MJT was supported by an AXA Postdoctoral Fellowship and NCBS campus fellowship. SM was supported by HFSP RGP0027/2012, JC Bose Fellowship, and a Wellcome Trust DBT-Alliance Margadarshi Fellowship (SM).

## Supplemental Information

**Fig. S1.**
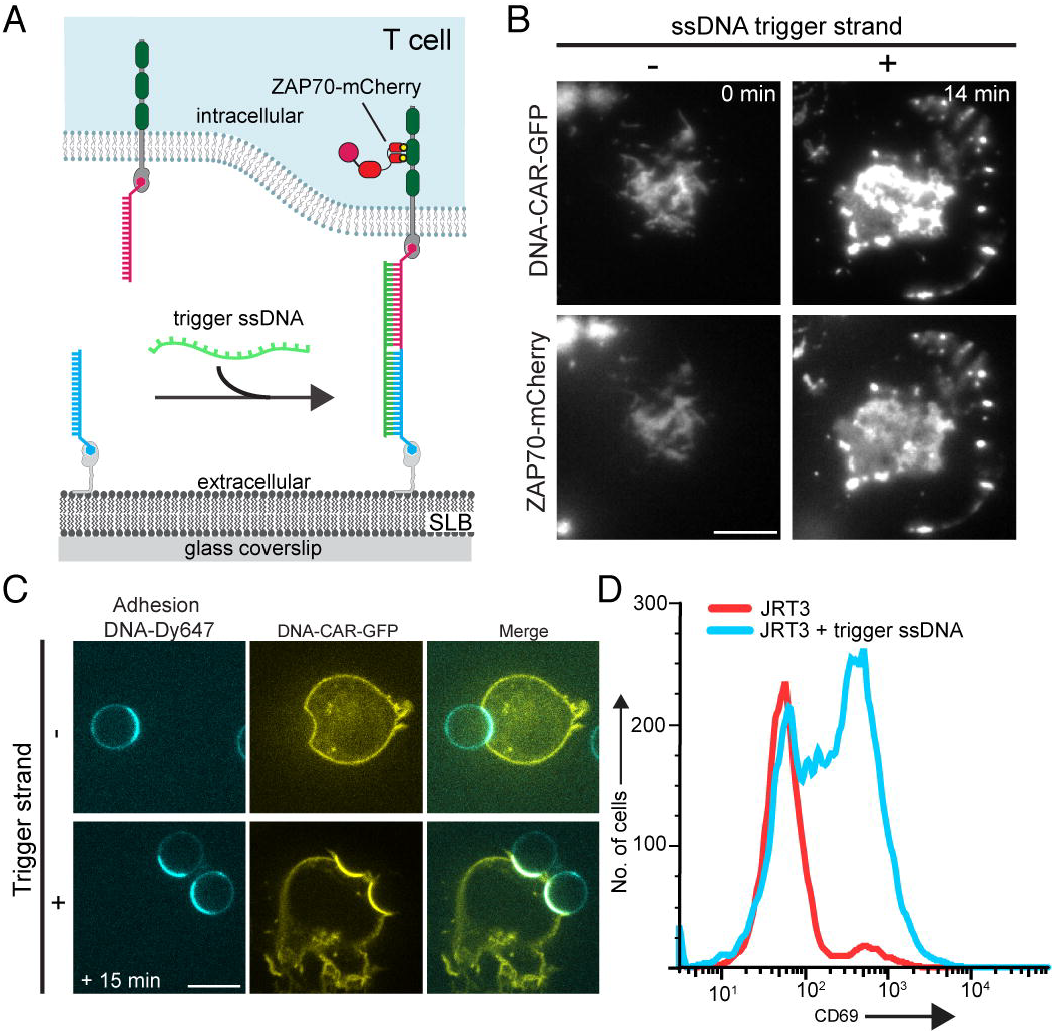
A triggerable DNA-CARζ system induces formation of ZAP70 positive microclusters and CD69 cell surface expression, related to Fig. 1. **A.** A schematic of a triggerable DNA-CARζ system with a 34 base pair overlap. **B.** Formation of ZAP70-mCherry positive microclusters and cell spreading after addition of DNA trigger strand which results in an overlap of 35 bp. Bar, 5 µm. Time after addition of the trigger DNA strand is shown. **C.** JRT3 cells expressing the DNA-CARζ-GFP were mixed with 7 µm silica microspheres with adsorbed supported lipid bilayers functionalized with a DNA adhesion strand (in this example, conjugated to His10-SNAPf-ybbr13-Dy647) and DNA ligand (conjugated to His10-CLIPf)). SLB were set up on silica microsphere to maintain cells in suspension and facilitate FACS analysis. Cells and beads were mixed in a 96 well plate before the addition of the DNA trigger strand. To check SLB formation on silica beads as well as DNA-CARζ activation, a portion of cells were plated onto poly-L-lysine coated coverslips and imaged by spinning disk confocal microscopy. Bilayer fluidity was checked by the enrichment of the His10-SNAPf-Dy647 at the cell-bead interface. 15 min after addition of trigger strand, the DNA-CARζ-GFP is enriched at the cell-microsphere interface. Scale bar, 10 µm. **D.** 4 hr after addition of trigger DNA strand, cells were stained for CD69 and analyzed by FACS. CD69 expression was upregulated on the cell surface for cells that were incubated with trigger strand (blue trace) compared with control cells (red trace).

**Fig. S2.**
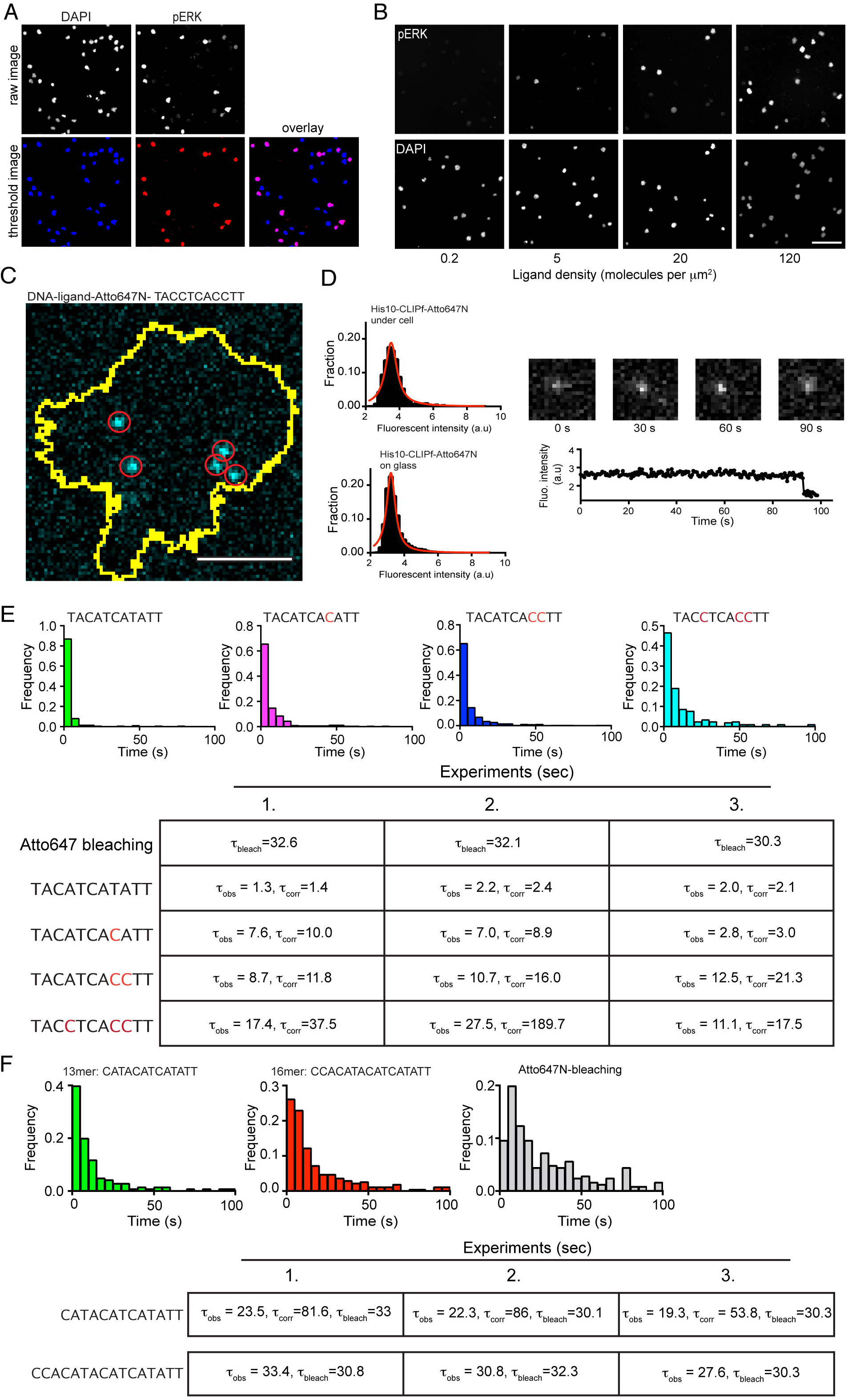
PhosphoErk quantification and measuring receptor-ligand dwell time, related to Fig. 2. **A.** A customized Cell Profiler analysis pipeline was used to process phosphoERK staining images and score JRT3 cells as phosphoERK positive or negative. First the DAPI channel raw images were segmented to identify the nucleus (shown as blue in the threshold image). Second the raw images of the phospoERK staining channeled were segmented using thresholding parameters set with control data sets (PMA stimulated cell, non-stimulated cells, and cells stained with secondary antibody only). Identified phosphoERK nuclei are shown as red in the thresholded image. Cells were scored phosphoERK positive when a DAPI nucleus overlapped with a segmented phosphoERK nucleus. A minimum threshold area was set for the nucleus as described in the Methods. In the overlay of the thresholded DAPI and phosphoERK images the nuclei scored positive appear magenta. Scale bar, 100 µm. **B.** Raw and threshold phosphoErk and DAPI images presented in Fig. 2A. Scale bar, 100 µm. **C.** JRT3 cells expressing DNA-CARζ were imaged using a two-camera TIRF microscope with simultaneous excitation of DNA-CARζ-GFP and DNA-ligand-Atto647N. A long 500 ms exposure in the Atto647N channel revealed single molecules (red circles) of DNA-ligands-Atto647N detected at the SLB-cell interface (yellow outline, obtained by segmentation using DNA-CARζ-GFP fluorescence). In contrast to bound DNA ligand, unbound DNA ligands diffuse more rapidly and appear as a unresolved fluorescent blur in images taken at 500 ms exposure. (See method of O’Donoghue et al.^1^) Scale bar, 2.5 µm. **D.** Top, mean fluorescent intensities of single receptor-bound DNA ligands versus single His10-CLIPf-ybbr labeled with Atto647N absorbed to glass and imaged with identical conditions. Bottom, representative intensity trace of a single receptor-bound DNA ligand labeled with Atto647N which is bound for 90 s, before disappearing by single step photobleaching/unbinding. **E-F.** Histograms of dwell times for DNA ligand and tables of individual dwell times (τ_obs_) and Atto647N bleaching measurements (τ_bl_). Panel c shows the 3 experiments that make up the average dwell times presented in Fig. 2d for the 11mer DNA ligands. Each experiment for the 11mer ligand represents measurement conducted on one experimental day. A single experiment consisted of single molecule dwell times measurement from >10 cells with 300-700 single receptor ligand binding events per DNA ligand analyzed. Histograms of individual dwell time measurements for the 13mer and 16mer DNA ligand are presented in panel (D). The τ_obs_ was corrected using the bleaching measurement performed during each experiment to obtain an estimate of average dwell time (τ_corr_) of receptor ligand unbinding from:

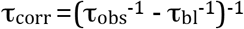 This method to estimate τ_corr_ is most accurate when τ_obs_ is considerably smaller that τ_bl_. Measurements of τ_obs_ that approach τ_bl_ have an inflated τ_corr_, as is the case the 11mer ligand with the greatest G/C content (AAGGTGAGGTA). In the case of the 16mer DNA ligand, τ_obs_ is equivalent to τ_bl_ making this correction method more prone to error.

**Fig. S3.**
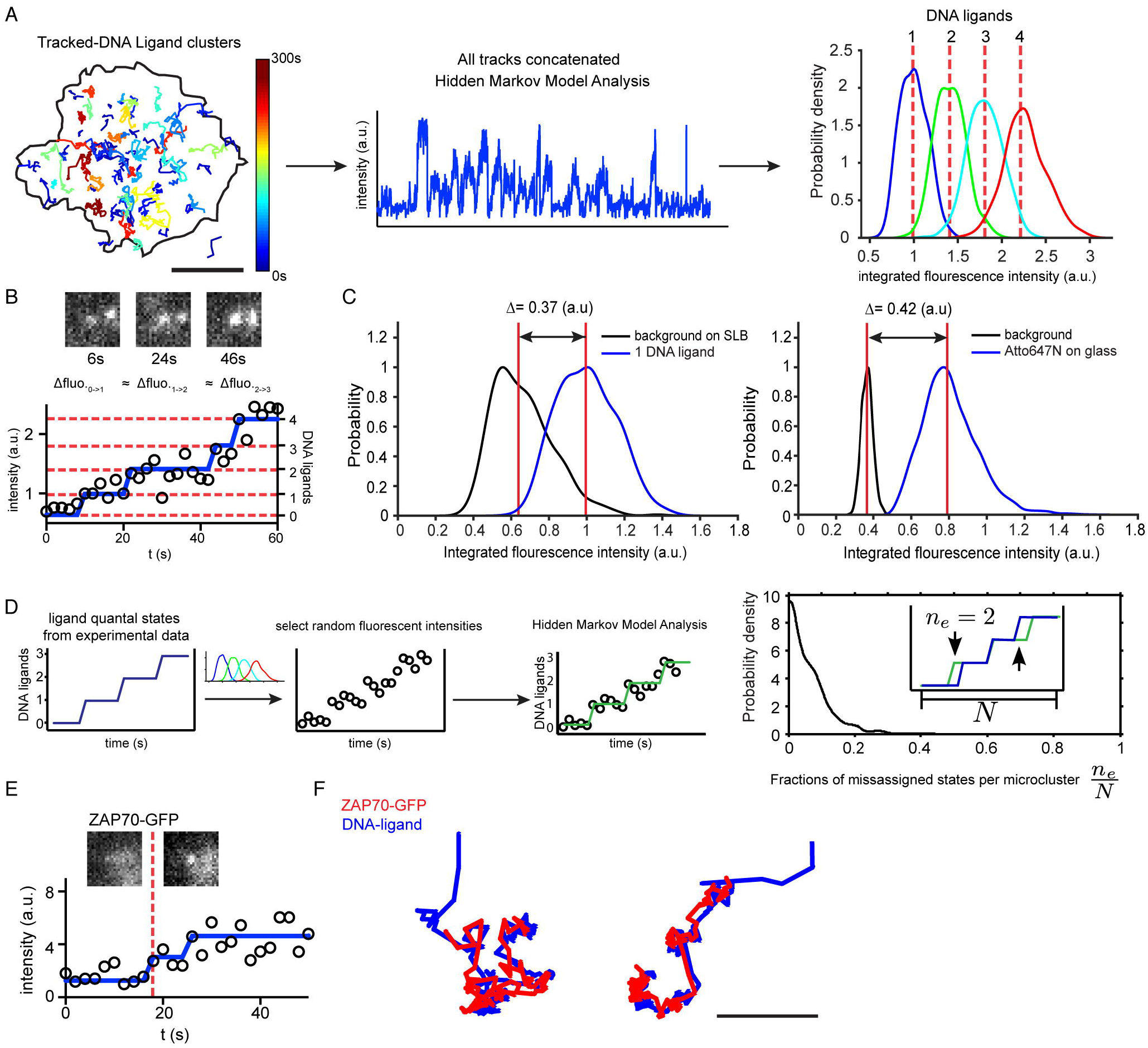
Analysis of receptor-ligand microcluster formation and ZAP70 recruitment, related to Fig. 3. **A.** We adapted a previously described^2^ Hidden Markov Model (HMM) analysis to identify the number of fluorescent ligands in each frame from the fluorescent intensity of tracked Atto647N ligand clusters. The intensity time-series of each tracked cluster was extracted from the coordinates generated by TrackMate. Shown is an example of 86 particle trajectories detected at the cell-SLB junction for the 16mer DNA ligand at ~1 molecule/µm^2^ (black outline, segmented cell boundary; microcluster trajectories colour coded by lifetime). Bar, 5 µm. The intensity time-series of all tracks were then concatenated, and the ensemble intensity time series was analyzed by a statistical maximum evidence approach, implemented with a variational Bayesian method. This analysis identified the rates of ligand addition and removal and the fluorescence distributions associated with n=1,2,3… bound ligands from the intensity data. We found the assigned distributions of fluorescent intensities are overlapping, as is expected from the highly fluctuating background noise generated by the fluorescent blur of the rapidly-diffusing unbound Atto647N-ligand on the camera detector. Nonetheless, each distribution shows a clear maximum and does not exhibit any structure indicative of misfitting (e.g. multiple peaks). Furthermore, the difference in the medians of each distribution peak is quantized, commensurate with the fluorescence intensity arising from a discrete numbers of fluorescent molecules. The identified rates and fluorescence distributions were used to reconstruct the time-series of ligand number for each cluster. This resulted in the identification of the step changes in the fluorescent signal that denoted new ligand binding events. **B.** Example DNA ligand intensity time series overlaid with HMM-assigned ligand number at each time point (blue trace). The step changes in the blue trace represent the binding of new ligands. The dashed red lines represent the median fluorescent intensity from n=1,2,3… fluorescent ligands (derived from the fluorescence intensity distributions shown in panel a). **C.** To ensure that the states identified by the HMM analysis corresponded to quantized intensity values characteristic of single molecules, we performed an identical analysis on data sets consisting of single Atto647N fluorophores stuck to glass slides. We then compared the fluorescent distributions of the n = 0 (i.e. background) and n = 1 fluorophore number with the distributions of the n=0 and n=1 fluorescent ligands found in the cell-bilayer data. We find that the difference in their medians is in good agreement (3.7 vs 4.2 fluorescent intensity units), confirming that the HMM analysis is reliably identifying states corresponding to discrete numbers of fluorescent molecules. We note that the HMM analysis was able to correctly identify states in both the stuck Atto647N dataset as well as the cell-based data-set despite the differences in noise between them (compare the distribution in the background on a SLB versus a glass slide). The background fluorescence intensity of the SLB was measured by sampling 5 time points previous to cluster appearance in a region of interest centered on the coordinate where the tracked microcluster appeared. The background distribution of the glass slide was obtained by randomly sampling regions in the image stack not occupied by single molecules. **D.** We assayed for the robustness of the HMM analysis to the broad fluorescence distributions by constructing a computer-generated data set. Time series of ligand binding and unbinding/bleaching were taken from an analyzed experimental dataset and converted into fluorescence values by sampling from the experimentally intensity distributions for n=1,2,3… DNA ligands shown in panel (a). This ensured that the noise and the transition rates between states in the computer-generated data set were reflective of experimentally collected data. The resultant time-series were analysed and the HMM reconstruction of state time series was compared to the original values. The error was quantified as the fraction of time-point in a time-series in which the HMM reconstruction misidentified the state. Plotting the distribution of error confirms that the algorithm is able to identify states with <10% error despite the noise present experimentally. **E.** We used the algorithm described in (a) conservatively to assay for presence or absence of ZAP70. Due to the noisy background signal associated with the cytosolic ZAP70-GFP, we checked the HMM assignment of the first ZAP70-GFP binding event by manual visual inspection to determine whether the detected step change in intensity represented recruitment. **F.** Representative DNA-ligand microclusters trajectories (blue) overlaid with the trajectory of an associated ZAP70-GFP spot (red). Positive ZAP70 recruitment was defined as a puncta of ZAP70-GFP that co-localized and co-migrated with an object in the ligand channel. By inspection of the data we found misidentified ZAP70 recruitment arose from two factors: 1) fluctuation in the Z-axis of the cell in the evanescent field, and 2) the presence of intracellular structures (likely endosomes) that were ZAP70-GFP positive that were prominent in some cells. These ZAP70-GFP punctae did not colocalise or move with ligand-bound receptors. Bar, 1 µm.

**Fig. S4.**
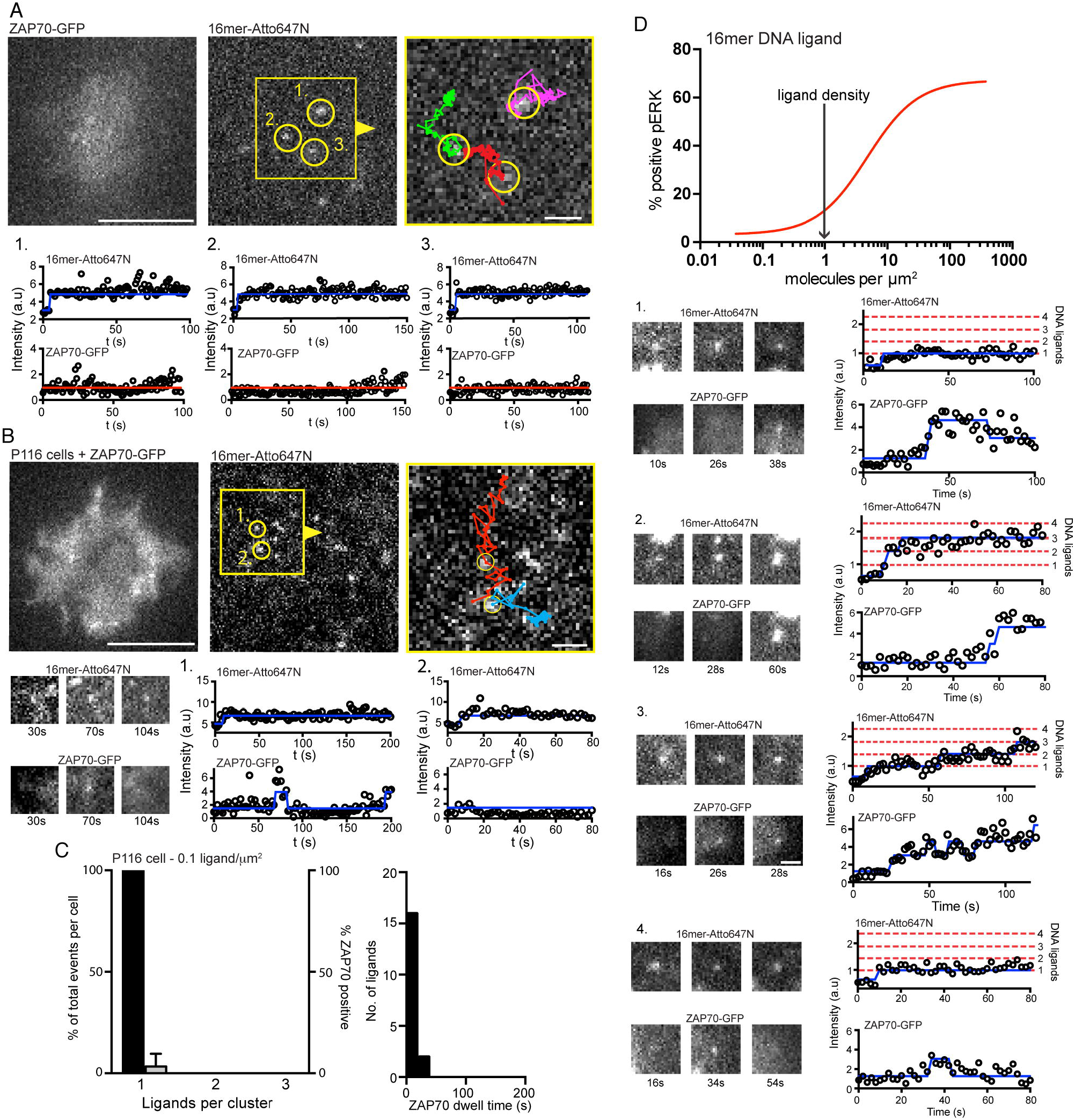
Gallery of single molecule binding events in JRT3 and P116 (ZAP70-negative Jurkat cell line) expressing DNA-CARz and ZAP70-GFP, related to Fig. 3. Single molecule imaging was performed at a density 0.1 16mer-DNA ligands/ µm^2^. **A.** TIRF images of 16mer DNA ligands labeled with Atto647N. Single bound ligands are marked by yellow circles in region of interest (red box and arrowhead). Bar, 5 µm. Right panel, region of interest overlaid with the tracked single molecule trajectories. ROI bar, 1 µm. Bottom panels, the fluorescent intensity time series for the single molecules of Atto647N labeled 16mer ligand and the corresponding ZAP70-GFP fluorescent intensity. Fluorescent intensity time series for ligand and ZAP70 channels analyzed as described in Fig. S3. **B.** TIRF images of a P116 cell expressing ZAP70-GFP and DNA-CARζ Bar, 5 µm. Region of interest (yellow box and arrowhead) on right overlaid with single molecule trajectories. ROI bar, 1 µm. Fluorescent intensity plots for bound ligands shown below images. **C.** Quantification of ZAP70 recruitment to single bound receptors in P116 cells. Shown are the number single bound ligand-receptor pairs (percent of total, black bars) and the percent of single bound receptor and clusters that recruited ZAP70-GFP (grey bars) in P116 Jurkat cells. ZAP70 recruitment was only observed in 2.2 ± 4% of single molecules events in P116 cells. Results are mean ± s.d. from 6 cells for P116 Jurkat cells. **D.** Quantification of ZAP70 dwell time at single bound ligands of 16mer DNA ligand in P116 cells (n=18). **E.** Gallery of examples showing formation of receptor-ligand microclusters and ZAP70 recruitment to a high affinity 16mer DNA ligand acquired by at a density of 1 ligand/µm^2^. The DNA ligand and ZAP70 fluorescent intensity time series were analyzed as described in Fig. S3. Receptor-ligand microclusters of 16mer DNA-ligand that formed in the initial 5 min of interacting with the SLB. Although uncommon, ZAP70 recruitment was observed at single molecules of bound 16mer ligand (examples 1-3). In some cases ZAP70 recruitment to single bound 16mer ligands was transient (example 4). Receptor-ligand microclusters consisting of 2 or more 16mer DNA ligands (examples 2) were more likely to recruit ZAP70. Bar, 1 µm.

**Fig. S5.**
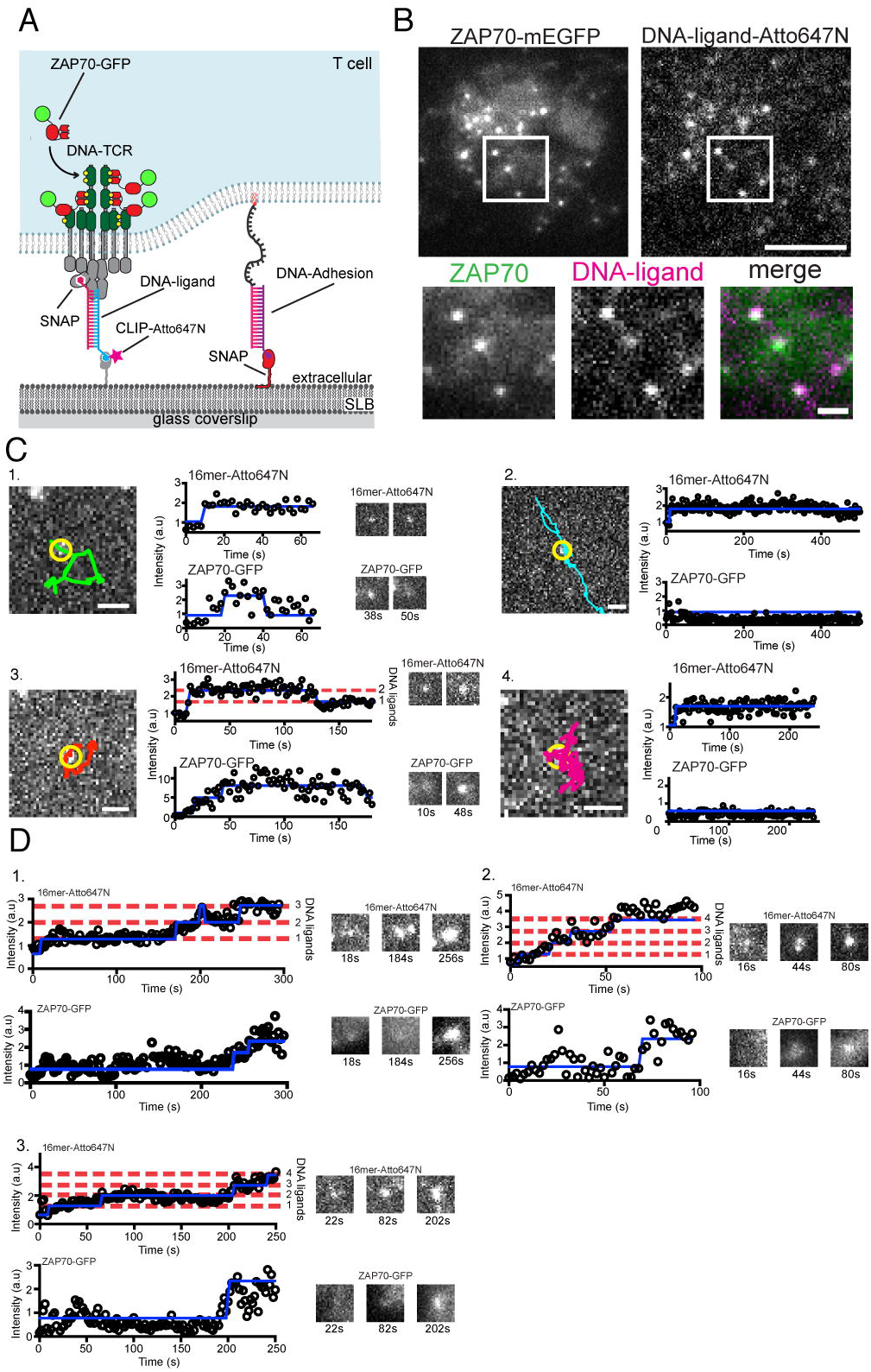
Recruitment of ZAP70 to DNA-CAR_TCR_ microclusters and galleries DNA-CAR_TCR_ binding events 0.1 and 1 16mer-DNA ligands/ µm^2^, related to Fig. 4. **A.** Schematic of the DNA-CARtcr. The SNAPf tag was fused to the extracellular domain of the TCRb and expressed in JRT3 cells (TCRb negative) to reconstitute the full TCR. **B.** TIRF microscopy images of a JRT3 Jurkat cell expressing ZAP70-GFP and DNA-CAR_TCR_ labeled with 16mer ssDNA after landing on a SLB with a complementary 16mer strand (120 molecules per mm^2^). Microclusters of ligand-receptor complexes formed within ~2 min and recruited ZAP70-GFP (inset). Scale bar, 5 µm; inset scale bar, 1 µm. **C.** Gallery of single molecule binding event at a density 0.1 16mer-DNA ligands/µm^2^. Left panels, TIRF images of 16mer DNA ligands labeled with Atto647N (yellow circle) and overlaid with the tracked single molecule trajectories. Right panels, the fluorescent intensity time series for the single molecules of Atto647N labeled 16mer ligand and the corresponding ZAP70-GFP fluorescent intensity. Fluorescent intensity time series for ligand and ZAP70 channels were analyzed as described in Fig. S3. Although uncommon, transient ZAP70 recruitment was observed at some single ligands bound to DNA-CAR_TCR_ (example 1). Example 3 shows a rare example of a small cluster consisting of two ligands forming, which also recruits ZAP70. Example 2 and 4, show more common single molecules DNA-CAR_TCR_ binding events that showed no discernable ZAP70 recruitment. Scale bar, 1 µm. **D.** Examples of DNA-CAR_TCR_ receptor-ligand microcluster formation and ZAP70 recruitment to a high affinity 16mer DNA ligand. Data shown were acquired by at a density of 1 ligand/µm^2^. The DNA ligand and ZAP70 fluorescent intensity time series were analyzed as described in Fig. S3. DNA-CAR_TCR_ receptor-ligand microclusters of 16mer DNA-ligand that formed in the initial 5 min of interacting with the SLB.

**Fig. S6.**
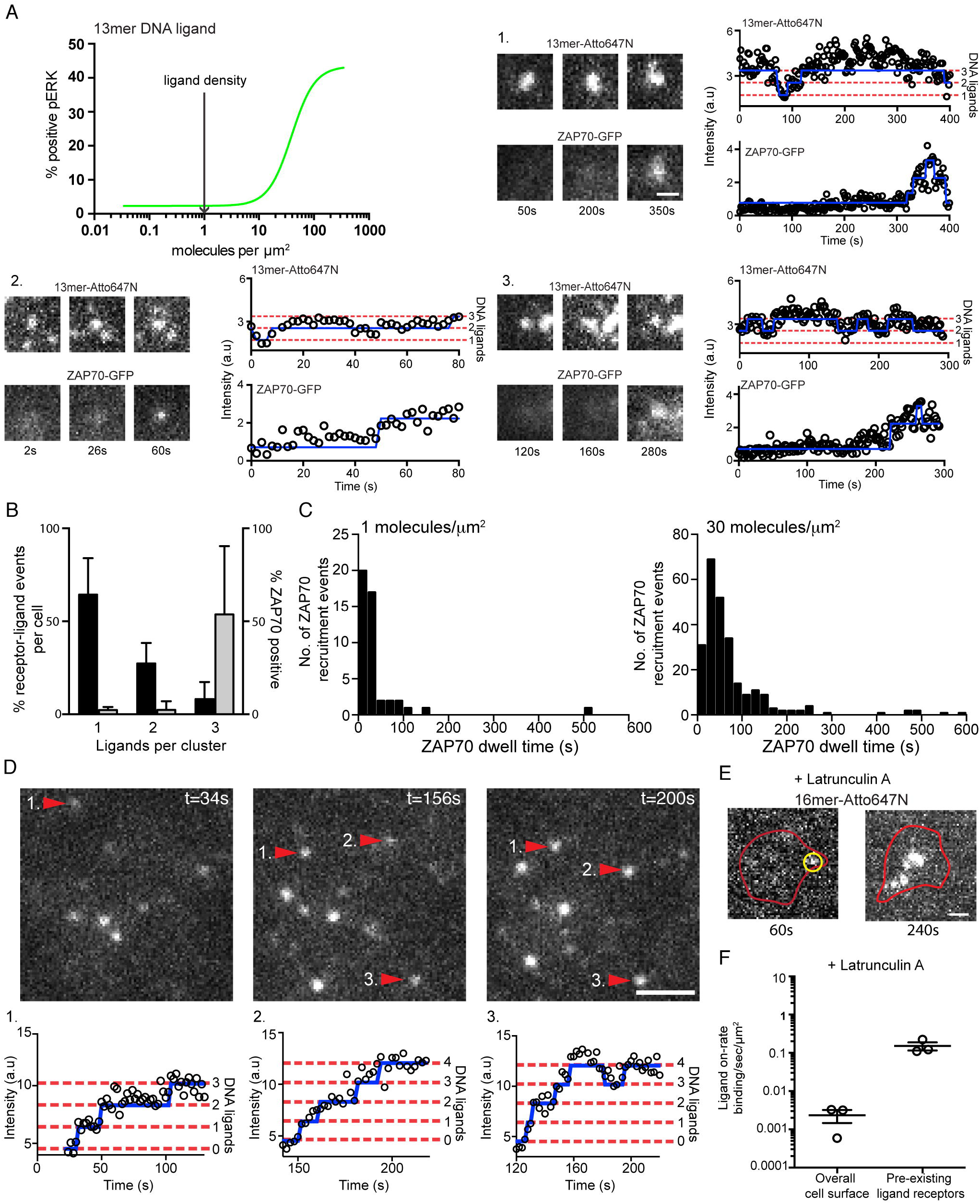
Receptor-ligand microclusters formation and ZAP70 recruitment to a low affinity 13mer DNA ligand, Related to Fig. 5. 13mer DNA ligand data shown acquired at a density of 1 ligand/µm^2^. The DNA ligand and ZAP70 fluorescent intensity time series were analyzed as described in Fig. S3. **A.** A gallery of examples showing the formation of small receptor-ligand microclusters of 13mer DNA ligand observed after cells had interacted SLBs for >15 min. **B.** Quantification of ZAP70 recruitment and dwell time to ligand-receptors of 13mer DNA ligand. Bar plot shows the number single bound ligand-receptor and clusters (percent of total binned, black bars) and quantification of (percent of total that recruit ZAP70, grey bars). Single molecule ligand binding events and receptor-ligand clusters of increasing ligand number that persisted for at least 30 s were scored for whether they recruited ZAP70-GFP. The initial point of ZAP70 recruitment was referenced to the number of ligands within the receptor-ligand spot. 53 ± 37% of receptor-ligand clusters containing 3 or more ligand recruit ZAP70 compared to 2 ± 1.7% of single bound ligands. Results are mean ± s.d. from n = 4 cells. **C.** The distribution of ZAP70 dwell times at receptor-ligand clusters at 1 ligand/pm^2^ (n=46 clusters from 4 cells) and 30 ligand/pm^2^ (N=276 cluster from 4 cells). Formation of DNA-CAR_TCR_ microclusters from single ligand-receptor binding events (D-E). **D.** A TIRF image of DNA-CAR_TCR_-bound 16mer DNA ligand organized into receptor-ligand clusters that grow by adjacent ligand binding events (red arrows numbered 1-3). Bar, 2.5 µm. Clusters grow by sequential addition of newly bound ligand; the blue line overlaid the fluorescent intensity represents detected step changes in fluorescence intensity (see methods and Fig. S3). Formation of DNA receptor ligand clusters in the presence of latrunculin A (E-F). **E.** TIRF images showing cluster formation nucleated from a single bound receptor (yellow circle) in the presence of latrunculin A (1 mM) with a 16mer at 1 ligand/µm^2^ (red outline show the cell outline obtained by segmentation using the ZAP70-GFP fluorescence). Bar, 1 µm. **F.** The rate of new ligand binding events near to an existing receptor-bound ligand (quantal increase in intensity in an existing diffraction-limited spot) or outside of these zones (measured by the sudden appearance of a new bound ligand in the contact area between the cell with the supported lipid bilayer) in cell treated with latrunculin A (1 mm). The ligand-receptor on-rate was expressed as the number of events per sec per µm^2^ membrane surface area (using 0.126 µm^2^ for a diffraction limited spot of a pre-existing ligand-receptor pair). Shown is the mean ± s.e.m. from 3 cells.

**Fig. S7.**
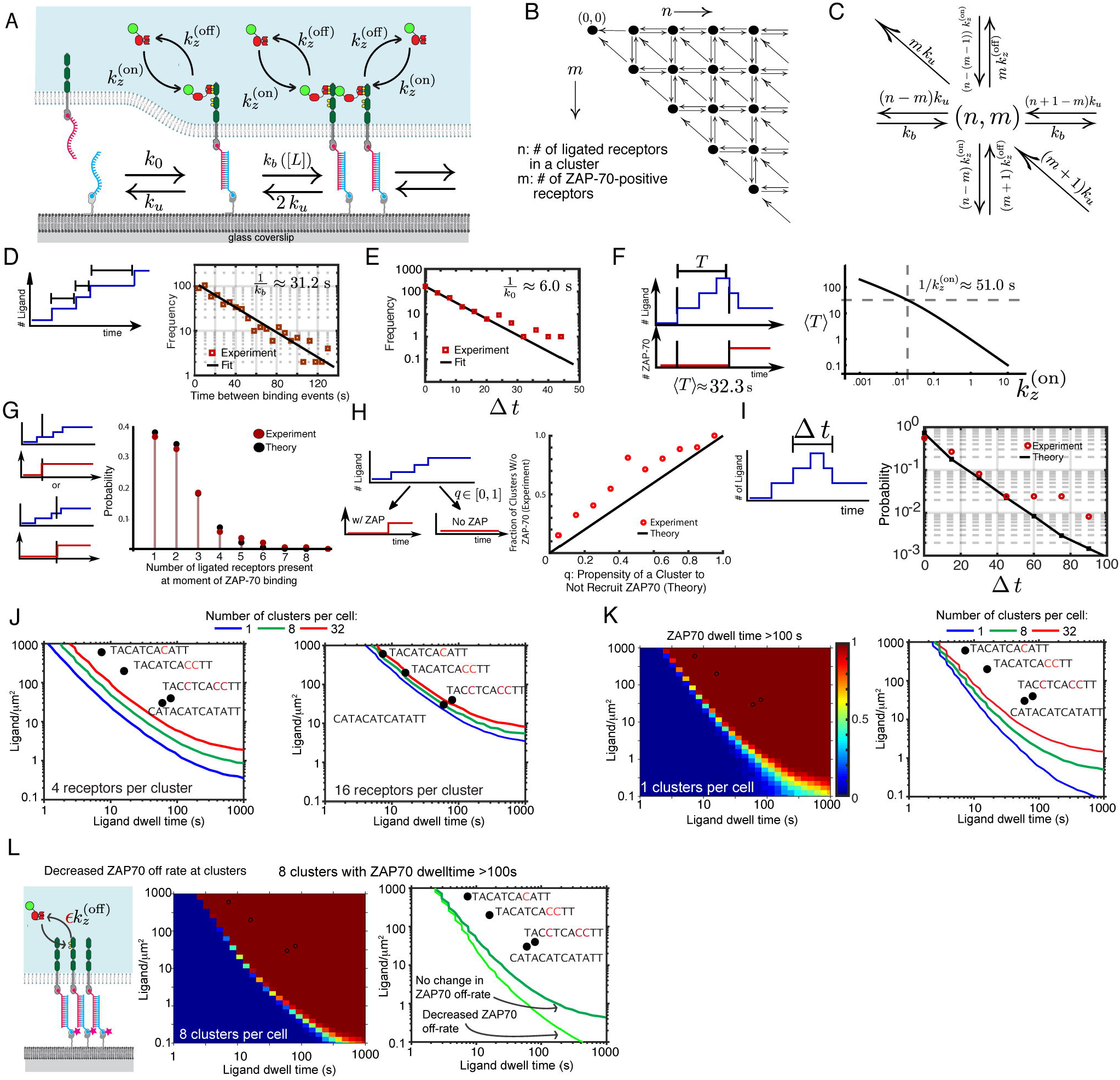
Construction of a theoretical signalling model. **A.** Schematic of the model, showing contact site formation (with rate *k*_0_), maturation into a cluster by additional ligand binding (with rate *k_b_*), ZAP70 binding and unbinding, and ligand unbinding. **B.** State space of an individual cluster. Each state is identified by the doublet of integers (*n,m*): the number of ligated receptors (n) and the number of those receptors that are ZAP70 positive (m). Note that *m ≤ n.* **C.** Possible stochastic transitions, and associated rates, from a representative state (*n, m*). Estimation of model parameters from experimental data:
**D.** Parameter *k_b_*: Histogram of waiting times between ligand binding at a cluster (red squares – 621 binding events over 434 tracked ligand clusters from 14 cells) with an exponential fit (black line) with rate constant *k_b_*. Left: schematic of measurement. **E.** Parameter 
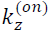
: Left: Schematic of measurement of the average time to first ZAP70 recruitment, <T>. Value obtained from 197 ZAP70 recruitment events from 14 cells. Right: <T> as a function of the ZAP70 on-rate, obtained from the model, with dashed lines indicating experimentally observed value (horizontal line) and corresponding rate (vertical line). **F.** Distribution of waiting times between appearances of ligand tracks, (381 waiting times from 14 cells). In red is an exponential fit with rate *k_Q_.* Validation of model against data:
**G.** Statistics of ZAP70 recruitment to clusters of ligated receptors, i.e. number of ligands present in cluster at the moment of the first ZAP70 recruitment event, for both theory (black) and experiment (red – from 197 ZAP70 recruitment events). **H.** Ligand tracks from experimental data (428, from 14 cells) binned by their propensity to not recruit ZAP70 (see text for details of calculation), compared with the fraction of each bin that do not, in fact, recruit ZAP70. Small but consistent deviation of data (red circles) from theoretical prediction (black line) indicates that less ZAP70 is seen experimentally than expected from the model. **I.** ‘Lifetime’ of a cluster, defined as n > 1 (see inset), for the 13mer ligand (black circles) against simulation (red squares and line). Simulation was performed with the bleaching-included ligand off-rate of 20 *s*^−1^. Note that the first four bins contain the majority of the experimental data (114 of 129 data points), with the last few bins containing 1 to 3 data points each. **J-L.** We quantitatively assessed the theoretical signalling model by performing stochastic simulations of cells interacting with a ligand functionalised supported lipid bilayer. These simulations were performed over a range of ligand densities (.01 to 1000 molecules per mm^2^) and ligand affinities (off-times of 1 to 1000 s) for a fixed time interval of 500 s. At each point in this parameter space we simulated 250 cells. The output of this simulation was time-series data of ligand binding, cluster formation and ZAP70 recruitment. We analysed this time series data for when clusters appeared with a particular number of ZAP70 positive receptors or ZAP70 association times. **J.** The number of ZAP70 receptors per cluster was used to define potential activation criteria. In example given the activation criteria is defined as the formation of a receptor-ligand cluster that have 4 or 16 or greater ligands. Coloured lines indicated the 20% contour from probability heat maps of when >20% of simulated cells forming the indicated number of clusters with a minimum of 4 or 16 ZAP70 positive receptors. To compare the predicted emergence of these features in the simulated data with experimental data, we plotted the ligand densities and ligand dwell time at which the indicated DNA ligands elicited a response of 20% phospho-Erk-positive cells (Fig. 2 and Fig. S2). **K.** Analysis of when clusters form with a ZAP70 association time of 100s. Left, heat map showing the probability of 1 cluster forming with a ZAP70 dwell time of 100s. Right, coloured lines indicated the 20% contour from probability heat maps of when >20% of simulated cells forming the indicated number of clusters with a ZAP70 dwell time of >100s. **L.** Schematic of a modified model in which the ZAP70 off rate is decreased at a cluster of ligated receptors (*∊<1*, see SI text for details) and heat map showing the results of simulation. Comparison of this model with a decreased dwell time of ZAP70 to the standard model, shows that decreased off rate of ZAP70 increases the probability of ZAP70 recruitment to high affinity ligand at lower ligand densities.

**Table S1.**
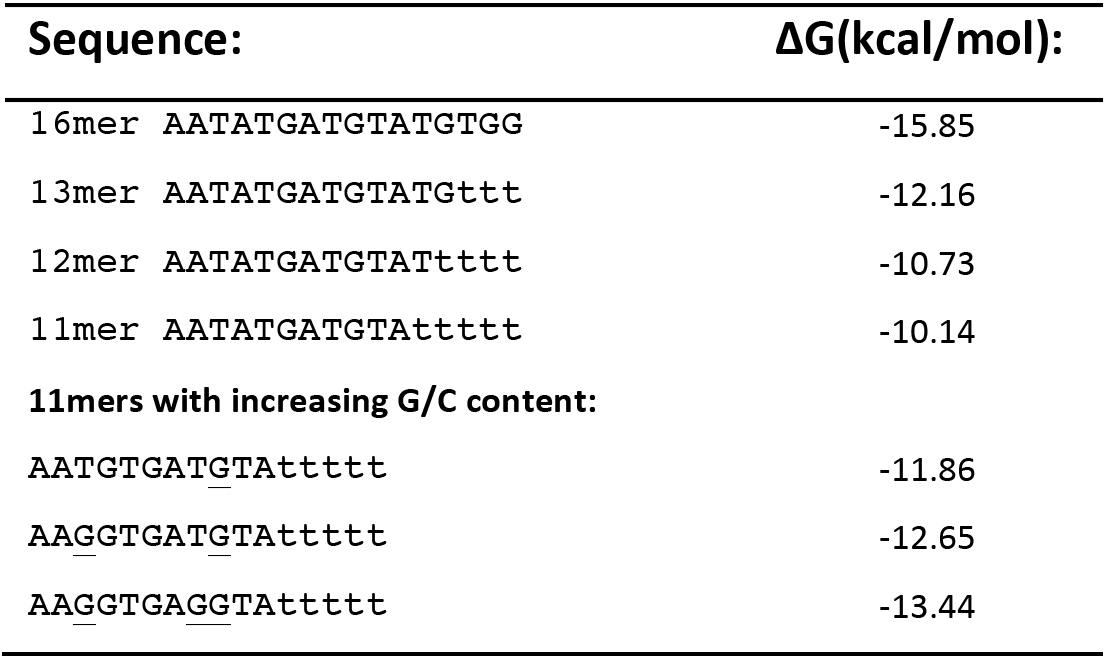
Computational analysis of ssDNA binding free energy. Calculated using nupack.org^3^ (accessed on 06/12/2014). Input parameter were a temperature of 37° C with 150 mM NaCl and 2.5 mM MgCl_2_.

## Supplemental Movie Captions

**Movie S1.** DNA-CARζ forms dynamic microclusters that recruitment ZAP70. This video shows a JRT3 expressing DNA-CARz and ZAP70-GFP (green channel) interacting with a supported lipid bilayer functionalized with 120 molecules/µm^2^ of 16mer DNA ligand-Atto647N (magenta channel). Video illustrates that the DNA-CARζ forms dynamic microclusters when engaged with a complementary DNA strand ligand, and DNA-CARζ microclusters recruit ZAP70 and move from the cell periphery to coalesce under the cell. Same cell as shown in Fig. 1B. Scale bar, 5 µm.

**Movie S2.** Calcium flux in JRT3 activated by an inducible DNA-CARt. JRT3 cells loaded with the Fura2 calcium dye were activated using an inducible DNA-CARζ, as described in Fig. 1D. Shown is the pseudo-color video of the Fura 2 emission ratio (340/380 nm excitation). The Fura 2 emission ration is constant until the addition of trigger strand at 200 seconds results cytosolic flux of calcium in a portion of cells at ~260 seconds. Scale bar, 5 µm.

**Movie S3.** Single molecule imaging of bound DNA ligands and ZAP70. Video showing single molecules of 16mer DNA ligand (magenta channel, bilayer density of 0.1 molecules/µm^2^) detected at the bilayer-cell interface of a JRT3 cell expressing DNA-CARζ and ZAP70-GFP (green channel). Single bound 16mer DNA ligands (labeled with atto647N) are detected as described in Fig. S2. Despite the long lifetime of the DNA-CARζ-16mer interaction, ZAP70-GFP recruitment is rarely seen at single ligand engaged receptors. Same cell as shown in Fig. S4A. Scale bar, 1 µm.

**Movie S4.** Monitoring the assembly of DNA-CARt-ligand microclusters and ZAP70 recruitment. This video shows a JRT3 expressing DNA-CARζ and ZAP70-GFP (green channel) interacting with a SLB functionalized 16mer DNA ligand labeled with atto647N (magenta channel, ligand density 1 molecules/µm^2^). At this ligand density the initial moment in the formation of small receptor-ligand microclusters can be detected. This movie shows that brighter receptor-ligand microclusters (consisting of 3 or more DNA ligands) more readily recruit ZAP70. Same cell as shown in Fig. 3D. Scale bar, 1 µm.

**Movie S5.** DNA-CAR_TCR_ forms microclusters that recruitment ZAP70. This video shows a JRT3 expressing DNA-CAR_TCR_ and ZAP70-GFP (green channel) interacting with a supported lipid bilayer functionalized with 100 molecules/pm^2^ of 16mer DNA ligand-Atto647N (magenta channel). Video illustrates that the DNA-CAR_TCR_ forms dynamic microclusters when engaged with a complementary DNA strand ligand, and DNA-CAR_TCR_ microclusters recruit ZAP70. Same cell shown in Fig. S5B. Scale bar, 5 µm.

**Movie S6.** DNA-CAR_TCR_ forms dynamic microclusters that move centripetally. This video shows a JRT3 expressing DNA-CAR_TCR_ (labeled with 16mer DNA strand receptor conjugated to a Cy3B fluorophore, red channel) interacting with a supported lipid bilayer functionalized with 120 molecules/µm^2^ of 16mer DNA ligand-Atto647N (cyan channel). Video illustrates that the DNA-CAR_TCR_ forms dynamic microclusters that move from the cell periphery to coalesce under the cell. Scale bar, 5 µm.

**Movie S7.** Single molecule imaging of bound DNA ligands and ZAP70 recruitment to the DNA-CAR_TCR_. Video showing single molecules of 16mer DNA ligand-Atto647N (magenta channel, bilayer density of 0.1 molecules/µm^2^) detected at the bilayer-cell interface of a JRT3 cell expressing DNA-CAR_TCR_ and ZAP70-GFP (green channel). Single bound 16mer DNA ligands are detected as described in Fig. S2. Scale bar, 1 µm.

**Movie S8.** A weaker 13mer DNA ligand and ZAP70 recruitment. Video showing a JRT3 expressing DNA-CARz and ZAP70-GFP (green channel) in in the initial minutes (<15 min after been after been applied to the bilayer) of interacting with a SLB functionalized 13mer DNA ligand labeled with atto647N (magenta channel, ligand density 1 molecules/µm2). In contrast to an equivalent density of the 16mer ligand, the majority of bound 13mer ligands are short lived and do not recruit ZAP70. Scale bar, 1 µm.

**Movie S9.** Receptor ligand microclusters of 13mer DNA ligand are more readily observed in cells after 15 mins. Video of a JRT3 cell after 15 min on a bilayer functionalized with 1 molecules/µm2 of 13mer DNA ligand (magenta channel). Microclusters of 13mer ligand were more readily observed after cell had been incubated with supported lipid bilayers for longer time period (>15 mins, Fig. 5A). ZAP70 recruitment (green channel) was almost exclusively observed a microclusters consisting of 3 or more DNA ligands. Examples of DNA microclusters and ZAP70 recruitment from this cell are shown in Fig. S6A. Scale bar, 1 µm.

**Movie S10.** Receptor ligand microcluster primarily assemble by the addition of newly bound ligand. Video of JRT3 expressing DNA-CARζ interacting with a bilayer functionalized with 1 molecules/µm2 of 16mer DNA ligand (same cell as shown in Fig. 6A). This video illustrates how microclusters primarily assemble by the unitary addition of newly bound ligand. The blue, red, and green overlays in the ligand channel (magenta) correspond to the three microclusters shown in Fig. 6A. The ZAP70 channel is shown in green. Scale bar, 1 µm.

**Movie S11.** Receptor ligand microcluster primarily assemble by the addition of newly bound ligand. Video of JRT3 expressing DNA-CAR_TCR_ interacting with a bilayer functionalized with 1 molecules/pm2 of 16mer DNA ligand (same cell as shown in Fig. S10). This video illustrates how microclusters primarily assemble by the unitary addition of newly bound ligand. The blue, red, and green overlays in the ligand channel (cyan) correspond to the three microclusters shown in Fig. S6C. In this video the DNA-CAR_TCR_ is labeled with a 16mer DNA strand receptor conjugated to a Cy3B (red channel). Scale bar, 2 µm.

## References

1. Straus, D. B. & Weiss, A. Genetic evidence for the involvement of the lck tyrosine kinase in signal transduction through the T cell antigen receptor. Cell 70, 585–593 (1992).

2. Reich, Z. et al. Ligand-specific oligomerization of T-cell receptor molecules. Nature 387, 617–620 (1997).

3. Shaw, A. S. & Dustin, M. L. Making the T cell receptor go the distance: a topological view of T cell activation. Immunity 6, 361–369 (1997).

4. Chan, A. C., Irving, B. A., Fraser, J. D. & Weiss, A. The zeta chain is associated with a tyrosine kinase and upon T-cell antigen receptor stimulation associates with ZAP-70, a 70-kDa tyrosine phosphoprotein. Proc. Natl. Acad. Sci. U.S.A. 88, 9166–9170 (1991).

5. Zhang, W., Sloan-Lancaster, J., Kitchen, J., Trible, R. P. & Samelson, L. E. LAT: the ZAP-70 tyrosine kinase substrate that links T cell receptor to cellular activation. Cell 92, 83–92 (1998).

6. Bunnell, S. C., Kapoor, V., Trible, R. P., Zhang, W. & Samelson, L. E. Dynamic actin polymerization drives T cell receptor-induced spreading: a role for the signal transduction adaptor LAT. Immunity 14, 315–329 (2001).

7. Cantrell, D. T cell antigen receptor signal transduction pathways. Annu. Rev. Immunol. 14, 259–274 (1996).

8. Brownlie, R. J. & Zamoyska, R. T cell receptor signalling networks: branched, diversified and bounded. Nature Publishing Group 13, 257–269 (2013).

9. Abraham, R. T. & Weiss, A. Jurkat T cells and development of the T-cell receptor signalling paradigm. Nat. Rev. Immunol. 4, 301–308 (2004).

10. Davis, M. M. & Bjorkman, P. J. T-cell antigen receptor genes and T-cell recognition. Nature 334, 395–402 (1988).

11. Gascoigne, N. R., Zal, T. & Alam, S. M. T-cell receptor binding kinetics in T-cell development and activation. Expert reviews in molecular medicine 2001, 1–17 (2001).

12. Davis, M. M. et al. Ligand recognition by alpha beta T cell receptors. Annu. Rev. Immunol. 16, 523–544 (1998).

13. Feinerman, O., Germain, R. N. & Altan-Bonnet, G. Quantitative challenges in understanding ligand discrimination by alphabeta T cells. Mol. Immunol. 45, 619–631 (2008).

14. Altan-Bonnet, G. & Germain, R. N. Modeling T cell antigen discrimination based on feedback control of digital ERK responses. PLoS Biol. 3, e356 (2005).

15. McKeithan, T. W. Kinetic proofreading in T-cell receptor signal transduction. Proc. Natl. Acad. Sci. U.S.A. 92, 5042–5046 (1995).

16. Rabinowitz, J. D. et al. Altered T cell receptor ligands trigger a subset of early T cell signals. Immunity 5, 125–135 (1996).

17. Klammt, C. et al. T cell receptor dwell times control the kinase activity of Zap70. Nat Immunol 16, 961–969 (2015).

18. Krogsgaard, M. et al. Evidence that structural rearrangements and/or flexibility during TCR binding can contribute to T cell activation. Mol. Cell 12, 1367–1378 (2003).

19. Schamel, W. W. A., Risueño, R. M., Minguet, S., Ortíz, A. R. & Alarcon, B. A conformation- and avidity-based proofreading mechanism for the TCR-CD3 complex. Trends in Immunology 27, 176–182 (2006).

20. Li, Q.-J. et al. CD4 enhances T cell sensitivity to antigen by coordinating Lck accumulation at the immunological synapse. Nat Immunol 5, 791–799 (2004).

21. O’Donoghue, G. P., Pielak, R. M., Smoligovets, A. A., Lin, J. J. & Groves, J. T. Direct single molecule measurement of TCR triggering by agonist pMHC in living primary T cells. Elife 2, e00778 (2013).

22. Wallace, V. A., Penninger, J. & Mak, T. W. CD4, CD8 and tyrosine kinases in thymic selection. Curr. Opin. Immunol. 5, 235–240 (1993).

23. Irving, B. A. & Weiss, A. The cytoplasmic domain of the T cell receptor zeta chain is sufficient to couple to receptor-associated signal transduction pathways. Cell 64, 891–901 (1991).

24. Eshhar, Z., Waks, T., Gross, G. & Schindler, D. G. Specific activation and targeting of cytotoxic lymphocytes through chimeric single chains consisting of antibody-binding domains and the gamma or zeta subunits of the immunoglobulin and T-cell receptors. Proc. Natl. Acad. Sci. U.S.A. 90, 720–724 (1993).

25. Sadelain, M., Brentjens, R. & Rivière, I. The basic principles of chimeric antigen receptor design. Cancer Discov 3, 388–398 (2013).

26. Gautier, A. et al. An Engineered Protein Tag for Multiprotein Labeling in Living Cells. Chemistry & Biology 15, 128–136 (2008).

27. Ohashi, P. S. et al. Reconstitution of an active surface T3/T-cell antigen receptor by DNA transfer. Nature 316, 606–609 (1985).

28. Grakoui, A. et al. The immunological synapse: a molecular machine controlling T cell activation. Science 285, 221–227 (1999).

29. Varma, R., Campi, G., Yokosuka, T., Saito, T. & Dustin, M. L. T cell receptor-proximal signals are sustained in peripheral microclusters and terminated in the central supramolecular activation cluster. Immunity 25, 117–127 (2006).

30. Yokosuka, T. et al. Newly generated T cell receptor microclusters initiate and sustain T cell activation by recruitment of Zap70 and SLP-76. Nat Immunol 6, 1253–1262 (2005).

31. Mor, A. et al. The lymphocyte function-associated antigen-1 receptor costimulates plasma membrane Ras via phospholipase D2. Nat. Cell Biol. 9, 713–719 (2007).

32. Selden, N. S. et al. Chemically programmed cell adhesion with membrane-anchored oligonucleotides. J. Am. Chem. Soc. 134, 765–768 (2012).

33. Weiss, A. & Littman, D. R. Signal transduction by lymphocyte antigen receptors. Cell 76, 263–274 (1994).

34. Smith-Garvin, J. E., Koretzky, G. A. & Jordan, M. S. T cell activation. Annu. Rev. Immunol. 27, 591–619 (2009).

35. Zadeh, J. N. et al. NUPACK: Analysis and design of nucleic acid systems. J Comput Chem 32, 170–173 (2011).

36. Kaizuka, Y., Douglass, A. D., Varma, R., Dustin, M. L. & Vale, R. D. Mechanisms for segregating T cell receptor and adhesion molecules during immunological synapse formation in Jurkat T cells. Proc. Natl. Acad. Sci. U.S.A. 104, 20296–20301 (2007).

37. Stefanová, I. et al. TCR ligand discrimination is enforced by competing ERK positive and SHP-1 negative feedback pathways. Nat Immunol 4, 248–254 (2003).

38. Kamentsky, L. et al. Improved structure, function and compatibility for CellProfiler: modular high-throughput image analysis software. Bioinformatics 27, 1179–1180 (2011).

39. Rabinowitz, J. D., Beeson, C., Lyons, D. S., Davis, M. M. & McConnell, H. M. Kinetic discrimination in T-cell activation. Proc. Natl. Acad. Sci. U.S.A. 93, 1401–1405 (1996).

40. Lord, G. M., Lechler, R. I. & George, A. J. A kinetic differentiation model for the action of altered TCR ligands. Immunol. Today 20, 33–39 (1999).

41. Chan, A. C., Iwashima, M., Turck, C. W. & Weiss, A. ZAP-70: a 70 kd protein-tyrosine kinase that associates with the TCR zeta chain. Cell 71, 649–662 (1992).

42. Williams, B. L. et al. Genetic evidence for differential coupling of Syk family kinases to the T-cell receptor: reconstitution studies in a ZAP-70-deficient Jurkat T-cell line. Mol. Cell. Biol. 18, 1388–1399 (1998).

43. Alam, S. M. et al. Qualitative and quantitative differences in T cell receptor binding of agonist and antagonist ligands. Immunity 10, 227–237 (1999).

44. Bunnell, S. C. et al. T cell receptor ligation induces the formation of dynamically regulated signaling assemblies. The Journal of Cell Biology 158, 1263–1275 (2002).

45. Manz, B. N., Jackson, B. L., Petit, R. S., Dustin, M. L. & Groves, J. T-cell triggering thresholds are modulated by the number of antigen within individual T-cell receptor clusters. 1–6 (2011). doi:10.1073/pnas.1018771108/-/DCSupplemental/pnas.1018771108_SI.pdf

46. Deindl, S. et al. Structural basis for the inhibition of tyrosine kinase activity of ZAP-70. Cell 129, 735–746 (2007).

47. Davis, S. J. & van der Merwe, P. A. The kinetic-segregation model: TCR triggering and beyond. Nat Immunol 7, 803–809 (2006).

48. James, J. R. & Vale, R. D. Biophysical mechanism of T-cell receptor triggering in a reconstituted system. Nature 487, 64–69 (2012).

49. Reth, M. Oligomeric antigen receptors: a new view on signaling for the selection of lymphocytes. Trends in Immunology 22, 356–360 (2001).

50. Dushek, O., Das, R. & Coombs, D. A role for rebinding in rapid and reliable T cell responses to antigen. PLoS Comput. Biol. 5, e1000578 (2009).

51. Schamel, W. W. A. et al. Coexistence of multivalent and monovalent TCRs explains high sensitivity and wide range of response. J. Exp. Med. 202, 493–503 (2005).

52. Lillemeier, B. F. et al. TCR and Lat are expressed on separate protein islands on T cell membranes and concatenate during activation. Nature Publishing Group 11, 90–96 (2010).

53. Dunne, P. D. et al. DySCo: quantitating associations of membrane proteins using two-color single-molecule tracking. Biophys. J. 97, L5–7 (2009).

54. James, J. R. et al. Single-molecule level analysis of the subunit composition of the T cell receptor on live T cells. Proc. Natl. Acad. Sci. U.S.A. 104, 17662–17667 (2007).

55. Qi, S. Y., Groves, J. T. & Chakraborty, A. K. Synaptic pattern formation during cellular recognition. Proc. Natl. Acad. Sci. U.S.A. 98, 6548–6553 (2001).

56. Hu, J., Lipowsky, R. & Weikl, T. R. Binding constants of membrane-anchored receptors and ligands depend strongly on the nanoscale roughness of membranes. Proc. Natl. Acad. Sci. U.S.A. 110, 15283–15288 (2013).

57. Krobath, H., Rózycki, B., Lipowsky, R. & Weikl, T. R. Binding cooperativity of membrane adhesion receptors. Soft Matter 5, 3354 (2009).

58. Huppa, J. B. et al. TCR-peptide-MHC interactions in situ show accelerated kinetics and increased affinity. Nature 463, 963–967 (2010).

59. Gillespie, D. T. Exact Stochastic Simulation of Coupled Chemical-Reactions. Journal of Physical Chemistry 81, 2340–2361 (1977).

60. Huang, J. et al. A single peptide-major histocompatibility complex ligand triggers digital cytokine secretion in CD4(+) T cells. Immunity 39, 846–857 (2013).

61. Liu, B., Chen, W., Evavold, B. D. & Zhu, C. Accumulation of dynamic catch bonds between TCR and agonist peptide-MHC triggers T cell signaling. Cell 157, 357–368 (2014).

62. Springer, T. A. Adhesion receptors of the immune system. Nature 346, 425–434 (1990).

63. Chen, L. & Flies, D. B. Molecular mechanisms of T cell co-stimulation and co-inhibition. Nature Publishing Group 13, 227–242 (2013).

64. Szymczak-Workman, A. L., Vignali, K. M. & Vignali, D. A. A. Design and Construction of 2A Peptide-Linked Multicistronic Vectors. Cold Spring Harbor Protocols 2012, pdb.ip067876-pdb.ip067876 (2012).

65. Yin, J., Lin, A. J., Golan, D. E. & Walsh, C. T. Site-specific protein labeling by Sfp phosphopantetheinyl transferase. Nat Protoc 1, 280–285 (2006).

66. Farlow, J. et al. Formation of targeted monovalent quantum dots by steric exclusion. Nature Methods 10, 1203–1205 (2013).

67. Zhang, D. Y., Turberfield, A. J., Yurke, B. & Winfree, E. Engineering entropy-driven reactions and networks catalyzed by DNA. Science 318, 1121–1125 (2007).

68. Douglass, E. F., Miller, C. J., Sparer, G., Shapiro, H. & Spiegel, D. A. A comprehensive mathematical model for three-body binding equilibria. J. Am. Chem. Soc. 135, 6092–6099 (2013).

69. Chan, Y.-H. M., van Lengerich, B. & Boxer, S. G. Effects of linker sequences on vesicle fusion mediated by lipid-anchored DNA oligonucleotides. Proc. Natl. Acad. Sci. U.S.A. 106, 979–984 (2009).

70. Schindelin, J. et al. Fiji: an open-source platform for biological-image analysis. Nature Methods 9, 676–682 (2012).

71. Edelstein, A., Amodaj, N., Hoover, K., Vale, R. & Stuurman, N. Computer control of microscopes using µManager. Curr Protoc Mol Biol Chapter 14, Unit14.20 (2010).

72. Hopfield, J. J. Kinetic proofreading: a new mechanism for reducing errors in biosynthetic processes requiring high specificity. Proc. Natl. Acad. Sci. U.S.A. 71, 4135–4139 (1974).

73. Ninio, J. Kinetic amplification of enzyme discrimination. Biochimie 57, 587–595 (1975).

## Supplemental References

1. O’Donoghue, G. P., Pielak, R. M., Smoligovets, A. A., Lin, J. J. & Groves, J. T. Direct single molecule measurement of TCR triggering by agonist pMHC in living primary T cells. Elife 2, e00778 (2013).

2. Bronson, J. E., Fei, J., Hofman, J. M., Gonzalez, R. L. & Wiggins, C. H. Learning rates and states from biophysical time series: a Bayesian approach to model selection and single-molecule FRET data. Biophys. J. 97, 3196–3205 (2009).

3. Zadeh, J. N. et al. NUPACK: Analysis and design of nucleic acid systems. J Comput Chem 32, 170–173 (2011).

